# Transcranial temporal interference stimulation increases slow waves during human NREM sleep

**DOI:** 10.64898/2026.11.21.768349

**Authors:** Ido Haber, Erin L. Schaeffer, Simone Bruno, Zhiwei Fan, Beril Mat, Larissa Albantakis, Stephanie G. Jones, Melanie Boly, Chiara Cirelli, Giulio Tononi

## Abstract

Slow waves are a hallmark of sleep homeostasis and a target for enhancing restorative sleep. Transcranial temporal interference stimulation (tTIS) can non-invasively deliver a low-frequency envelope to deep targets and has been shown to increase spectral slow-wave activity during sleep. It remains untested whether tTIS also alters discrete slow waves, how their cortical expression changes, and whether the field each brain receives relates to the response. During non-rapid eye movement sleep, 32 participants received 1 Hz tTIS and 11 a carrier-only control in 3-min episodes targeting the left insula/ventromedial prefrontal cortex. Combining high-density electroencephalography (EEG), source reconstruction and individualized field modeling, we found that tTIS increased slow-wave density by about 20% during and 30% after stimulation, across shallow and deep waves, with limited morphology change. The carrier-only control showed no detected density change. In source space, each wave involved more cortex in both arms, while wave origins did not change. A dose-response analysis showed that tTIS participants who received a stronger modeled field shifted further toward N3 sleep during stimulation, a relation not seen in the control. These results support tTIS as a candidate EEG-compatible tool for modulating slow waves and motivate individualized field modeling to interpret its effects.

## Introduction

Sleep is a fundamental, evolutionarily conserved state that is essential for healthy brain function across the lifespan, supporting cognition, learning and memory, emotional regulation, and metabolic and immune health ^1–3^. Its importance is underscored by the consequences of its disruption. Insufficient or fragmented sleep impairs attention and memory and disturbs mood, and disordered sleep is a core feature of neurodegenerative and psychiatric disease ^2,4^. Sleep is not uniform but cycles between non-rapid eye movement (NREM) and rapid eye movement (REM) stages, and it is during deep NREM sleep that the brain expresses its most prominent low-frequency rhythms ^5^. These NREM slow rhythms are widely regarded as a central signature of restorative sleep and have been linked to synaptic homeostasis, cellular restoration, and memory consolidation ^6,7^. Their physiological importance makes them an attractive target for interventions that aim to enhance the restorative action of sleep itself.

One way to act on these rhythms is with non-invasive brain stimulation. Conventional transcranial electrical stimulation (tES), including transcranial direct- and alternating-current stimulation (tDCS and tACS), can shift cortical excitability and entrain ongoing oscillations by passing weak currents between scalp electrodes ^8,9^. However, the delivered current is largest in the superficial cortex beneath the electrodes and cannot be concentrated at a deeper target without also stimulating the overlying tissue ^10^. Transcranial temporal interference stimulation (tTIS) addresses this limitation by applying two high-frequency currents that differ by an offset, so that where the fields overlap, their superposition forms an amplitude-modulated envelope, while the carriers themselves remain in the kilohertz range ^11^. Because the envelope, rather than the carrier, is thought to be the primary, physiologically relevant signal, the locus of effective stimulation can in principle be optimized toward deeper structures than conventional tES ^11,12^. A growing body of human work has since used tTIS to produce measurable, target-specific physiological and behavioral effects ^13,14^.

Several properties make tTIS particularly well suited to combining stimulation with electroencephalogram (EEG) sleep recording. Because the kilohertz carriers sit far above the typical frequencies of interest on EEG, physiological data can be recorded at the same time as the stimulation, provided that dedicated filtering attenuates the carriers and recording-chain nonlinearities are controlled ^11,15,16^. Critically for sleep, perception thresholds rise steeply with carrier frequency, so kilohertz stimulation can be delivered with little or no perceptible sensation provided the amplitude is titrated per participant ^17^, reducing the chance that participants are awakened or become aware of stimulation.

Within NREM sleep, the dominant low-frequency activity is carried by slow waves, large-amplitude events occurring within the broader 0.5–4 Hz delta band, whose spectral power is termed slow-wave activity (SWA). These slow rhythms reflect near-synchronous transitions of cortical neurons between depolarized ON states and hyperpolarized OFF states ^18–21^. Within the delta band, slow waves are discrete events whose density, amplitude, slope, and spatial extent change systematically with sleep pressure across the night and serve as interpretable markers of homeostatic drive and NREM sleep depth ^20,22^. Decades of work show that these events can be modulated by magnetic, auditory, and electrical stimulation, though the effect depends strongly on state, timing, and modality ^3^. Transcranial magnetic stimulation (TMS) pulses elicit individual slow waves during NREM sleep but not during wakefulness ^23^, and auditory stimulation enhances ongoing slow oscillations only when timed to their up-state ^24^. With tES, slow-oscillatory tDCS and 0.5 Hz tACS enhanced slow oscillations and deep sleep ^25,26^, but these effects have not replicated consistently, and intracranial recordings found no entrainment of sleep rhythms by 0.75–1 Hz tACS at tolerable intensities ^27^.

tTIS has recently been applied during sleep. Schaeffer et al. (2026) reported that tTIS with 15 kHz carriers and a 1 Hz offset increased spectral SWA during NREM sleep overnight and that the effect persisted after stimulation ^16^. In that study, participants did not report sensation associated with stimulation during the night, and stimulation did not increase arousals, awakenings, or wake after sleep onset relative to baseline nights ^16^. Bruno et al. (2026) showed that tTIS with 15 kHz carriers and a 10 Hz offset targeting the thalamus could increase spindle-range sigma activity during a daytime nap ^28^. These findings establish that tTIS can modulate sleep oscillations, but several questions remain open. Can tTIS modulate slow waves at the level of discrete detected events rather than band-limited power? Also, can such changes be projected to and quantified in source space? Lastly, does an individual’s physiological response depend on the received dose? The present study extends previous work focused on spectral findings ^16^ to these three levels—discrete slow-wave events, source-space analysis, and a dose-response relationship. To address these questions, we compared tTIS (1 Hz beat) against a carrier-only control during NREM sleep, with stimulation targeted to the left insula/ventromedial prefrontal cortex, a key node in the distributed cortical network expressing sleep slow waves ^19,29^. Each stimulation episode was divided into consecutive pre-stimulation, stimulation and post-stimulation intervals (Fig. 1C). Slow waves were detected from high-density EEG and analyzed in both sensor and source space (Fig. 2).

**Fig. 1.**
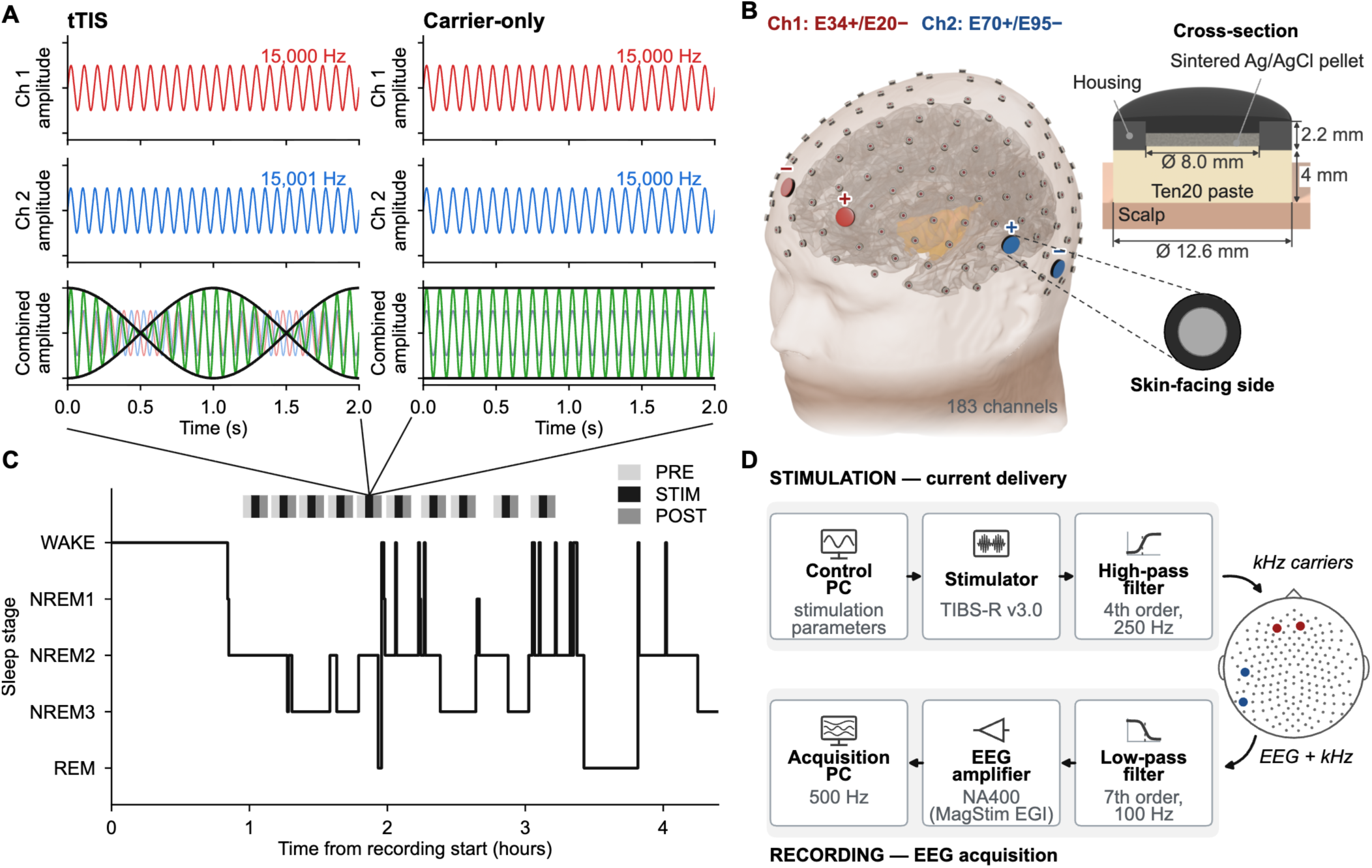
tTIS stimulation overview. (A) Stimulation waveforms for the tTIS and carrier-only control conditions: the two high-frequency carrier currents (red, 15,000 Hz; blue, 15,001 Hz in tTIS and 15,000 Hz in the carrier-only control) and their superposition (green). In the tTIS condition, the 1 Hz carrier offset produces an amplitude-modulated envelope (black), whereas the matched carriers of the carrier-only control produce no envelope. (B) Template head model (Ernie) showing the high-density EEG net and the four Ag/AgCl stimulation electrodes—channel 1 (E34+/E20−, red) and channel 2 (E70+/E95−, blue)—on the scalp surface, the modeled left-insula target (yellow), and the stimulation electrode, a sintered Ag/AgCl pellet (8.0 mm diameter) in a ring housing (12.6 mm outer diameter, 2.2 mm thick), shown in cross-section on the scalp with Ten20 conductive paste between the pellet and the skin (top; the paste is drawn at the 4 mm of the field model) and from its skin-facing side (bottom; the dashed lines join it to one of the stimulation electrodes on the head). (C) Hypnogram of one stimulation night, with the stimulation episodes marked above as matched PRE (light gray), STIM (black), and POST (mid gray) windows confined to NREM sleep across the early night. (D) Acquisition signal chain: stimulation current is generated by a control PC and TIBS-R v3.0 stimulator, high-pass filtered (4th order, 250 Hz), and delivered to the head as kHz carriers carrying the 1 Hz beat; the recorded EEG is low-pass filtered (7th order, 100 Hz) to attenuate the kHz carriers and digitized by the NA400 (MagStim EGI) amplifier and acquisition PC at 500 Hz.

**Fig. 2.**
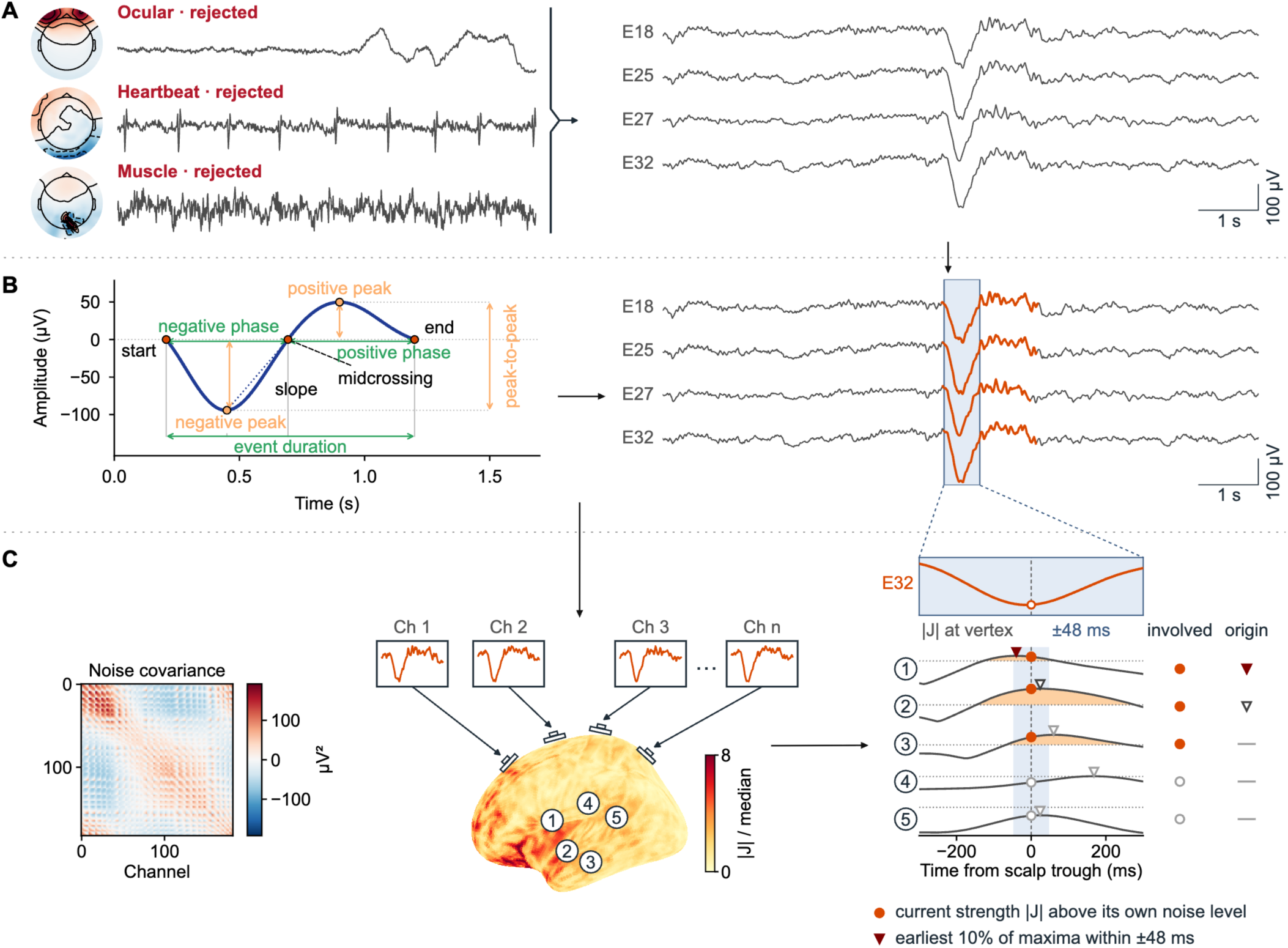
Signal processing pipeline. (A) ICA artifact removal: one AMICA decomposition per participant (maximum 1,000 iterations), manual inspection and rejection of artifact components (shown, one ocular, one heartbeat and one muscle component of one participant). Right, the cleaned EEG (average reference). (B) Slow-wave detection: waves are detected per channel with YASA on the 0.3–1.5 Hz band-passed signal. Left, YASA algorithm illustration. Right, the same EEG with the waves detected on each channel highlighted. (C) Source reconstruction: waves detected on several channels within 1 s are merged into one, and the source current at the merged wave’s negative peak is estimated with sLORETA on the participant’s own forward model and morphed to fsaverage5. The panel follows the wave highlighted in B. Left, the noise covariance of this participant. Middle, the wave of B and its source current at the negative peak on the left hemisphere, as the current strength |J| divided by each vertex’s median at random PRE times; the numbers mark five example vertices. Right, how the wave is scored. A vertex is involved (filled circle) when its current at the scalp trough exceeds its own 95th percentile at random times (dotted). Among the vertices whose maximum of relative current (triangle) falls within ±48 ms of the trough (band) and reaches 25% of the wave’s largest value, the earliest 10% form the origin (filled triangle); a dash marks a vertex that does not enter the origin ranking.

First, we hypothesized that tTIS would increase slow-wave density relative to the immediately preceding pre-stimulation period, an increase that would linger into the post-stimulation period, in line with the spectral SWA increase reported previously ^16^. Second, because slow waves are traveling, distributed cortical events, rather than the output of a single generator ^19,29^, we predicted that their expression would increase globally. We further asked whether any such increase reflected a shift in where waves originate and how much cortex they involve. Third, we hypothesized that inter-individual differences in modeled modulation depth (TI_max_), estimated from individualized head models, would positively correlate with the magnitude of the physiological response, and would not follow the strength of the high-frequency carrier field^10,30–32^.

## Methods

### Study Design and Demographics

Data from 43 participants (32 tTIS and 11 carrier-only; ages 18–49; Table 1) were analyzed from the STRENGTHEN study ^16^. Recruitment began with an online survey, followed by a telephone eligibility screen, and an in-person screening. Eligible participants were 18–50 years old. Major exclusion criteria were recent psychiatric illness, neurological disease (including obstructive sleep apnea), a history of seizures or significant head trauma, metal implants or other MRI contraindications, medications that lower seizure threshold, recent substance use disorder, and regular night-shift work; the full criteria are listed in the trial registration ^33^. All participants were medically healthy, with no acute illness, no diagnosed chronic illness or sleep disorders, and no medications expected to influence the outcome. Individuals completed a baseline magnetic resonance imaging (MRI) scan and a stimulation-free baseline sleep night. All participants gave written informed consent, and the study was approved by the University of Wisconsin–Madison Institutional Review Board (protocol 2023-1473). The STRENGTHEN study is registered at ClinicalTrials.gov (NCT06267521).

**Table 1.** Participant characteristics by arm.

|  | **tTIS (n = 32)** | **Carrier-only (n = 11)** | **p** |
|---|---|---|---|
| Age (years), mean ± SD [range] | 31.2 ± 9.7 [18–48] | 25.5 ± 8.2 [18–49] | 0.077 |
| Female, n (%) | 22 (68.8%) | 9 (81.8%) | 0.70 |

*Note. Age was compared between arms with a Mann–Whitney U test and sex with Fisher's exact test.*

Participants received tTIS or a carrier-only control during NREM sleep. Arm assignment was parallel and non-randomized, made after the baseline assessment with matching on age and sex, and participants were blind to their arm ^16^. Each stimulation episode was treated as a local time series of three consecutive intervals, pre-stimulation (PRE), stimulation (STIM), and post-stimulation (POST), with PRE and POST each equal in duration to STIM. The main contrasts compared STIM with PRE and POST with PRE so that each stimulation episode was anchored to its immediately preceding sleep state.

### tTIS Protocol

tTIS was delivered during NREM sleep using two high-frequency sinusoidal carriers at 15,000 Hz and 15,001 Hz; their superposition produced a 1 Hz amplitude-modulated interference envelope, selected to match the approximate frequency of endogenous sleep slow waves (Fig. 1A). Stimulation was generated by a TI Brain Stimulator for Research (TIBS-R v3.0, TI Solutions AG, Switzerland) and delivered through four sintered Ag/AgCl ring electrodes (8 mm pellet in a 12.6 mm housing, 2.2 mm thick) embedded in the high-density EEG net in place of recording electrodes and affixed with conductive paste (Ten20). Electrode impedances were monitored throughout the night and electrodes were adjusted if impedance rose substantially (above approximately 6.5 kΩ). A standardized two-channel montage of the GSN-HydroCel 256-channel net (EGI) was used for all participants, with the same polarity in both arms, channel 1 E34 (+) to E20 (−) and channel 2 E70 (+) to E95 (−) (Fig. 1B). In three tTIS participants, channel 2 used the adjacent electrode E94 in place of E95 because of an electrode placement and impedance issue. Every in silico model used the electrodes actually used, and the modeled fields of these three participants fell within the range of the others.

The montage was designed to target the left insula/ventromedial prefrontal cortex. Each channel was driven at ∼5 mA zero-to-peak. Across participants, the measured delivered current was approximately 5.2 (channel 1) and 5.1 mA (channel 2) zero-to-peak, and electrode impedance during stimulation was ∼4.8 kΩ (Fig. S1 and Table S1). Before sleep onset, with the participant awake, short test protocols (15 s with 15-s ramps) were delivered at increasing amplitudes up to 5 mA zero-to-peak to ensure no scalp sensations to stimulation ^16^. After sleep onset, a technician initiated stimulation once stable NREM sleep (at least 3 minutes) was identified online from the high-density EEG and polysomnography. Each episode comprised a 3-minute stimulation window with 15-second ramp-up and ramp-down periods (both included in the STIM interval). Episodes were separated by at least 6 minutes, and approximately ten episodes were delivered per night, in most cases confined to NREM sleep within the first four hours after sleep onset (Fig. 1C).

### EEG Preparation, ICA, and Interpolation

High-density EEG was acquired with a GSN-HydroCel 256-channel Ag/AgCl net and NA400 amplifier (MagStim EGI) sampled at 500 Hz, together with polysomnographic channels (electrooculography, submental electromyography, and electrocardiography). Dedicated hardware filters enabled simultaneous stimulation and recording. A high-pass filter (4th order, 250 Hz cutoff) on the stimulator output attenuated DC offsets and nonlinear low-frequency products ^15,34^ and a low-pass filter (7th order, 100 Hz cutoff) on the EEG amplifier input attenuated the kHz carriers to reduce the risk of amplifier saturation (Fig. 1D, Fig. S2). An optical trigger logged stimulation onset and offset for offline alignment.

Sleep was staged manually in 30-second epochs by a registered technician following American Academy of Sleep Medicine criteria ^35^, using the electrooculography, electromyography, and six high-density EEG channels approximating F3, F4, C3, C4, O1, and O2. Recordings were then aligned to stimulation triggers, sleep-stage annotations, channel metadata, and PRE/STIM/POST labels, and the continuous data were restricted to NREM stage 2 and 3 epochs and concatenated into one analysis-ready recording per participant. Preprocessing included zero-phase finite impulse response (FIR) band-pass filtering (0.5–40 Hz) and removal of non-EEG and visually identified bad channels. Semi-automated artifact rejection then removed epochs with excessive low- or high-frequency power and channels whose artifacts spanned most of the recording, followed by manual review. Acquisition, hardware filtering, and staging follow Schaeffer et al. ^16^.

Further artifact cleaning used independent component analysis (ICA) with AMICA (v1.5) in EEGLAB (v2025.1.0) under MATLAB (R2023b) ^36,37^. ICA was run with one model, a maximum of 1,000 iterations, and one processing thread per dataset. Components reflecting ocular, muscle and heartbeat activity were rejected, as were, in a few participants, components carrying a residual 1 Hz stimulation artifact not fully removed by filtering alone ^38^. A median of 13 components (range 2–36) of a median of 193 (range 141–227) was rejected per participant; the retained components preserved a median of 96.9% of channel-summed variance per 10-s segment (range 63.3–99.7%). After component rejection and review, channels excluded during cleaning were interpolated back to a common 183-channel working montage. A median of 38 of the 183 channels per participant (20.8%; range 7.1–41.0%) were interpolated as a common montage is required for the channel-wise cluster-permutation statistics. Interpolation used spherical splines implemented in MNE-Python (v1.12.1) ^39,40^.

### Slow-Wave Detection and Sensor-Space Metrics

Slow waves were detected independently on each channel of the cleaned NREM EEG, re-referenced to the common average of the 183 retained channels, after resampling to 100 Hz and band-pass filtering at 0.3–1.5 Hz, using the Yet Another Spindle Algorithm (YASA, v0.7.0) *sw_detect* function ^22,41,42^. Accepted events had negative-half-wave duration between 0.3 and 1.5 s and positive-half-wave duration between 0.1 and 1.0 s, and their negative-peak (40–200 microvolts), positive-peak (10–150 microvolts) and peak-to-peak (75–350 microvolts) amplitudes had to lie within YASA’s default bounds scaled to each participant (Fig. 2B; Fig. S3). Because scalp slow-wave amplitude differs severalfold between individuals and declines with age ^22^, fixed microvolt bounds would detect proportionally more waves in high-amplitude participants. Every amplitude bound was therefore multiplied by a participant scale factor k_i_ = s_i_/s_ref_, where s_i_ is the median over channels of each channel’s median trough depth across all PRE negative half-waves lasting 0.3–1.5 s, and s_ref_ = 10.94 µV is the cohort median of s_i_ (k_i_ 0.46–2.14, median 1.00). The scale was set from PRE alone, before any stimulation, and the same bounds applied to a participant’s PRE, STIM and POST intervals (Fig. S3). Sensor-space metrics were slow-wave density, negative-peak amplitude, positive-peak amplitude, peak-to-peak amplitude, slope and frequency (Table S3). Slow-wave density was defined as the number of detected slow waves in an interval divided by the usable NREM duration.

### Source Reconstruction: Slow-Wave Origin and Involvement

Source-space analyses used the detected slow-wave events, after collapsing channel-level detections of the same wave so that each slow wave was counted once ^19^. A detection joined a wave when it fell within 1.0 s of the wave’s first detection and its channel had not yet detected in that wave, and a second detection on the same channel opened a new wave. The most-negative channel served as the wave’s reference channel. For each subject, forward solutions were rebuilt from the MRI-derived individualized head models ^43^ and electrode positions. Forward models were then projected to the fsaverage5 cortical surface ^44^ for group analysis ^40^. The inverse used the same 183-channel montage as the sensor-space analysis. For source analysis, the cleaned EEG was band-pass filtered at 0.3–1.5 Hz, resampled to 250 Hz, and given an average-reference projection. Noise covariance was estimated from pre-stimulation segments, excluding every channel-level slow-wave detection on any channel from 0.5 s before its start to 1.0 s after its positive peak, with the remaining wave-free data divided into 2-second epochs. The covariance was then regularized by Ledoit–Wolf shrinkage ^45,46^. An inverse operator was then built with MNE minimum-norm tools using sLORETA, signal-to-noise ratio (SNR) = 3, lambda2 = 1/9, loose orientation constraint = 0.2, and depth weighting = 0.8 ^47^.

From each reconstructed wave we derived two complementary descriptions of its cortical expression, involvement and origin, both referenced to each vertex’s own resting current. Let J_v_(t) be the magnitude of the sLORETA current estimate at vertex v and t_w_ the negative peak of wave w on its reference channel. The resting current was sampled at random PRE times without a slow wave (one candidate per PRE wave, 1.5–3 s from its negative peak, kept when at least 1 s from any detected wave), and θ_v_ and m_v_ are the 95th percentile and the median of J_v_ over these times.

Involvement measures how much cortex a wave recruited above its resting level. Origin marks where the cortex peaked first, following the principle that a traveling slow wave begins where the cortex peaks earliest ^19,29^. For vertex v in wave w they are

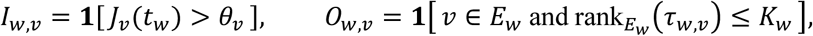

where 1[·] is 1 when the condition holds and 0 otherwise. For origin, τ_w,v_ is the latency of the local maximum of the relative current J_v_(t)/m_v_ nearest t_w_ within ±48 ms, E_w_ is the set of vertices whose such maximum exceeds 25% of the largest relative current of any vertex in that window, rank orders the latencies in E_w_ from the earliest, and K_w_ = max(1, round(0.10 |E_w_|)) keeps the earliest 10%, with vertices tied at the cutoff sharing the remaining places. Each probability map (P) is the mean of I_w,v_ or O_w,v_ over a participant’s waves in an interval (PRE, STIM or POST). The maps were then smoothed over fsaverage5 vertex adjacency.

### Individualized Electric Field Modeling

Individualized electric field modeling was performed through the TI-Toolbox (v3.1.0) ^48^, which provides the study-level interface for montage definition, individualized head-model handling, tTIS field computation, surface projection, and region-wise analysis ^32^. From each model, the modulation depth was derived at every cortical location. For carrier fields E1 and E2 and a unit direction n, the modulation depth is

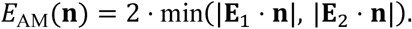

TI_normal_ evaluates this quantity along the midcortical (central) surface normal n_c_, whereas TI_max_ is its maximum over all directions, obtained with the closed-form solution of Grossman et al. ^11^:

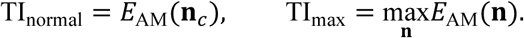

The high-frequency carrier dose HF_max_ is the peak carrier field ^49^. In tTIS, the relative phase of the two carriers cycles once per beat, so the peak is reached when the two fields add in alignment or in opposition. In the carrier-only control, the two phase-locked 15,000 Hz carriers are in phase, so they add coherently into one fixed field ^50^, with E1 and E2 signed by each channel’s hardware polarity:

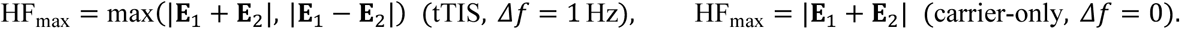

No envelope was delivered in the carrier-only arm, so its TI_max_ and TI_normal_ were computed from its two simulated carrier fields as the modulation depth they would have produced with a 1 Hz offset, a montage-matched placebo dose that carries each participant’s anatomy on the same scale.

Head models were segmented with CHARM (SimNIBS v4.5.0) from each participant’s T1- and T2-weighted structural MRI ^32,43^, acquired on a 3 T head-only scanner (MAGNUS, GE Healthcare) with a 32-channel head coil (T1-weighted MP-RAGE, repetition time 2000 ms, echo time 3 ms, inversion time 1100 ms, flip angle 8°; T2-weighted CUBE, repetition time 2500 ms, echo time 90 ms; both 0.8 mm isotropic). Stimulation electrodes were modeled as 8 mm diameter discs with a 4 mm gel layer (1.0 S/m) and a 2 mm electrode layer (29.4 S/m), driven with each participant’s measured channel currents (mean plateau current over all eligible logged protocols). All field summaries are modeled estimates computed with isotropic tissue conductivities (white matter 0.126, gray matter 0.275, cerebrospinal fluid 1.654, bone 0.01 [compact 0.008, spongy 0.025], scalp 0.465, eyes 0.5, blood 0.6, muscle 0.16 S/m) ^43,51^. Because different field measures yield different values and highlight different cortex ^52^, we prespecified TI_max_, TI_normal_, and HF_max_ as three surface-level field summaries. In the terminology of recent consensus work on dose-response relationships in transcranial stimulation ^53^, these summaries are individualized estimates of the received dose, that is, model-based reconstructions of the tissue-level field derived by combining the montage, electrode geometry, and delivered current with each participant’s anatomy and assumed tissue properties.

### Statistical Analysis

All map-level inference used cluster-based permutation tests (Fig. S4) ^54^, which control the family-wise error rate across channels or vertices within each map; all tests were two-sided. Every map used a rank statistic at each channel or vertex, namely the Wilcoxon signed-rank statistic for changes within a group, the Mann–Whitney U statistic for change scores between groups, and the Spearman correlation for field-response maps. Candidate clusters joined adjacent channels (high-density EEG layout) or vertices (cortical surface) with element-wise p < 0.05 (two-tailed); positive and negative clusters were formed separately. Cluster mass was compared at α = 0.05 with the maximum-cluster-mass null from 5,000 permutations (within-group sign flips, between-group label permutations or, for field-response maps, reassignment of responses among participants).

The false discovery rate was controlled with the Benjamini–Hochberg procedure ^55^ at q < 0.05, separately within each family of tests. Effect sizes were rank-biserial correlations (r) ^56,57^ of participant-level means over the significant cluster (the smallest-p cluster where a map had two), with subject-level bootstrap 95% confidence intervals (CI; 5,000 resamples)^58^.

Within-night trends were assessed by fitting, for each participant, an ordinary least-squares slope of the channel-averaged STIM-minus-PRE or POST-minus-PRE change on episode number and testing the slopes against zero with a two-sided Wilcoxon signed-rank test.

A post hoc analysis related dose to the shift toward N3 sleep. For each stimulation, the N3 share of an interval was the time scored N3 divided by the time scored N2 or N3, and its change from PRE was averaged over the participant’s stimulations. This change was correlated with the whole-cortex mean dose (Spearman, bootstrap 95% CI, 10,000 resamples), and the difference in ρ between groups was tested by permuting group labels (10,000 permutations). To locate where the local field tracked this change, participants’ dose at each vertex was correlated with their change in N3 share (Spearman) and tested with the cluster procedure above ^59^, with a cluster-forming threshold of p < 0.01. Clusters were labeled with the Desikan–Killiany atlas ^60^.

Correlations inside selected clusters and their adjustment for age and sex are descriptive and uncorrected. Leave-one-subject-out (LOSO) sensitivity results are listed in Table S5.

## Results

### tTIS Increases Slow-Wave Density During and After Stimulation

During stimulation, tTIS participants showed a slow-wave density increase across 174 channels (q = 0.0036; r = 0.85, 95% CI [0.64, 0.98]; cluster-mean increase of 0.29 waves/min (+20.0%); Fig. 3A). This increase lingered into the post-stimulation period across 176 channels (q = 0.0036; r = 0.83, 95% CI [0.59, 0.98]; cluster-mean increase of 0.44 waves/min (+30.5%); Fig. 3B). Carrier-only control participants showed no density change in either contrast (q = 0.95 for STIM/PRE and 0.53 for POST/PRE). The between-group tTIS-versus-carrier-only contrast formed a 30-channel cluster during stimulation (cluster p = 0.030) that did not survive correction across the 36 sensor maps (q = 0.18), and none after stimulation (q = 0.53). PRE density was lower in the carrier-only arm (channel-mean median 0.85 versus 1.31 waves/min in tTIS; Mann–Whitney U p = 0.044). Arm assignment was not randomized but matched on age and sex ^16^, so a baseline difference of this kind was not excluded by design.

**Fig. 3.**
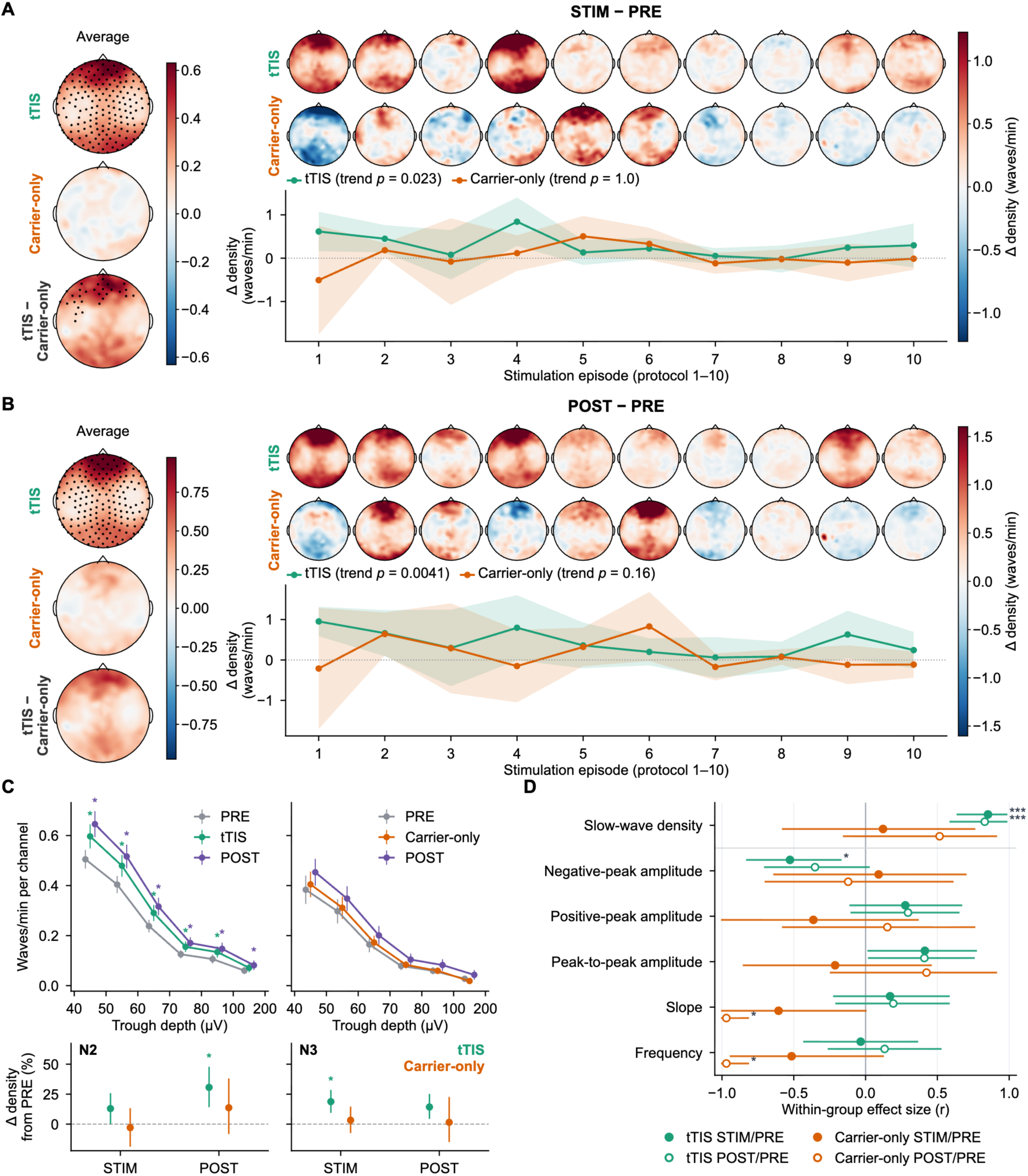
Sensor-space slow-wave density and morphology. (A) STIM − PRE and (B) POST − PRE density contrasts. Left, pooled density-change topomaps (averaged across all stimulation episodes) for tTIS, carrier-only and tTIS − carrier-only, with permutation-significant channels (corrected p < 0.05 within the map) marked as dots; the 30 tTIS − carrier-only channels during stimulation did not survive correction across the sensor maps (Table S3). Right, per-episode density-change topomaps (tTIS row over carrier-only row) above the within-night density trend across the first ten stimulation episodes. Points show the group-mean change at each episode (± subject-level 95% CI); p is the within-night trend test (Wilcoxon signed-rank test of participants’ slopes across episodes), given for each group in the legend. Color encodes the mean density change (waves/min). (C) Top, density over the tTIS STIM/PRE density-significant channels in six trough-depth bins for PRE, STIM and POST (participant mean ± SEM). Bottom, change in density from PRE within N2 and within N3 separately, as a percentage of each participant’s own PRE density in that stage (mean and participant-level bootstrap 95% CI). Asterisks mark a segment versus PRE (Wilcoxon signed-rank on the paired densities, Benjamini– Hochberg q < 0.05, separately over the 24 bin tests and the 8 stage tests). (D) Within-group effect sizes (r, subject-level bootstrap 95% CI) for slow-wave density and five morphology metrics, for tTIS and carrier-only (filled = STIM/PRE, open = POST/PRE), over the tTIS STIM/PRE density-significant channels. Asterisks mark the Wilcoxon signed-rank test after Benjamini–Hochberg correction across the 24 tests of the panel (* q < 0.05, ** q < 0.01, *** q < 0.001).

The increase in wave density spanned shallow and deep waves alike. In tTIS, density rose by 18–26% during and by 28– 39% after stimulation in each of six trough-depth bins (q < 0.05 for every bin except 100–200 µV during stimulation, q = 0.12), with no change in any carrier-only bin (Fig. 3C). Broken down by sleep stage, tTIS density rose within both N2 and N3 (by 13% to 31% of each participant’s own PRE density in that stage), significantly within N2 after stimulation (+31%) and within N3 during stimulation (+19%; q = 0.0096 for both), whereas carrier-only density did not change (−3% to +14%; all q ≥ 0.87).

The effect was strongest for wave density, with more limited changes in wave morphology (Fig. 3D; Table S3). Morphology clusters formed only for slow-wave frequency after stimulation, which rose in 13 channels in tTIS and fell in 11 channels in the carrier-only control, each significant within its map but not after correction across the sensor maps (q = 0.16 and 0.15; Fig. S6), and which differed between groups in 25 channels (q = 0.0096). Within the tTIS density cluster, paired within-participant changes showed deeper negative peaks during stimulation (r = −0.53, q = 0.040) and a smaller peak-to-peak amplitude increase during and after stimulation that did not survive correction (r = 0.41 for both, uncorrected p = 0.041 and 0.043, q = 0.15), without comparable changes in positive-peak amplitude, slope or frequency. In the carrier-only control, slope and frequency decreased in these channels after stimulation (r = −0.97 for both, q = 0.012; Fig. 3D). These changes were consistent across participants but small. In tTIS the negative peak deepened by a mean of 0.5 µV (1.0% of each participant’s PRE value) and peak-to-peak amplitude rose by 0.7 and 0.6 µV (0.8%), and in the carrier-only control slope fell by 4.2 µV/s (2.0%) and frequency by 0.014 Hz (1.8%), against a density increase of 20–30% in tTIS.

Importantly, the density increase was not uniform across the night. Treating each stimulation episode as a time point, the within-tTIS density gain declined across episodes for both contrasts (per-participant slope versus episode: STIM/PRE p = 0.023, r = −0.46; POST/PRE p = 0.0041, r = −0.57), with no corresponding trend in the carrier-only control (p = 1.0 and 0.16; Fig. 3A,B). No wave-morphology trend reached significance in tTIS (all p ≥ 0.32), and in the carrier-only control only negative-peak amplitude during stimulation showed one (p = 0.014, r = −0.85; Fig. S6; Table S3). Episode order and elapsed time were nearly collinear, so the declining density response could not be reliably attributed to either time of night or cumulative stimulation (Fig. S7).

### Slow Waves Involve More Cortex in Both Arms

Per wave, the probability that a slow wave involved a given vertex rose in tTIS during stimulation across two significant clusters totaling 36.6% of cortical-surface vertices (q = 0.014; r = 0.72) and after stimulation across two significant clusters totaling 60.6% (q = 0.014; r = 0.77; Fig. 4). It also rose in the carrier-only control, during stimulation in one right inferior parietal and lateral occipital cluster (12.8%; q = 0.032) and after stimulation in two clusters (38.9%; q = 0.014), and the groups did not differ (q = 0.11 for STIM/PRE and 0.15 for POST/PRE). Origin probability did not change within either group or between them (all q ≥ 0.065; Fig. S8; Table S4).

**Fig. 4.**
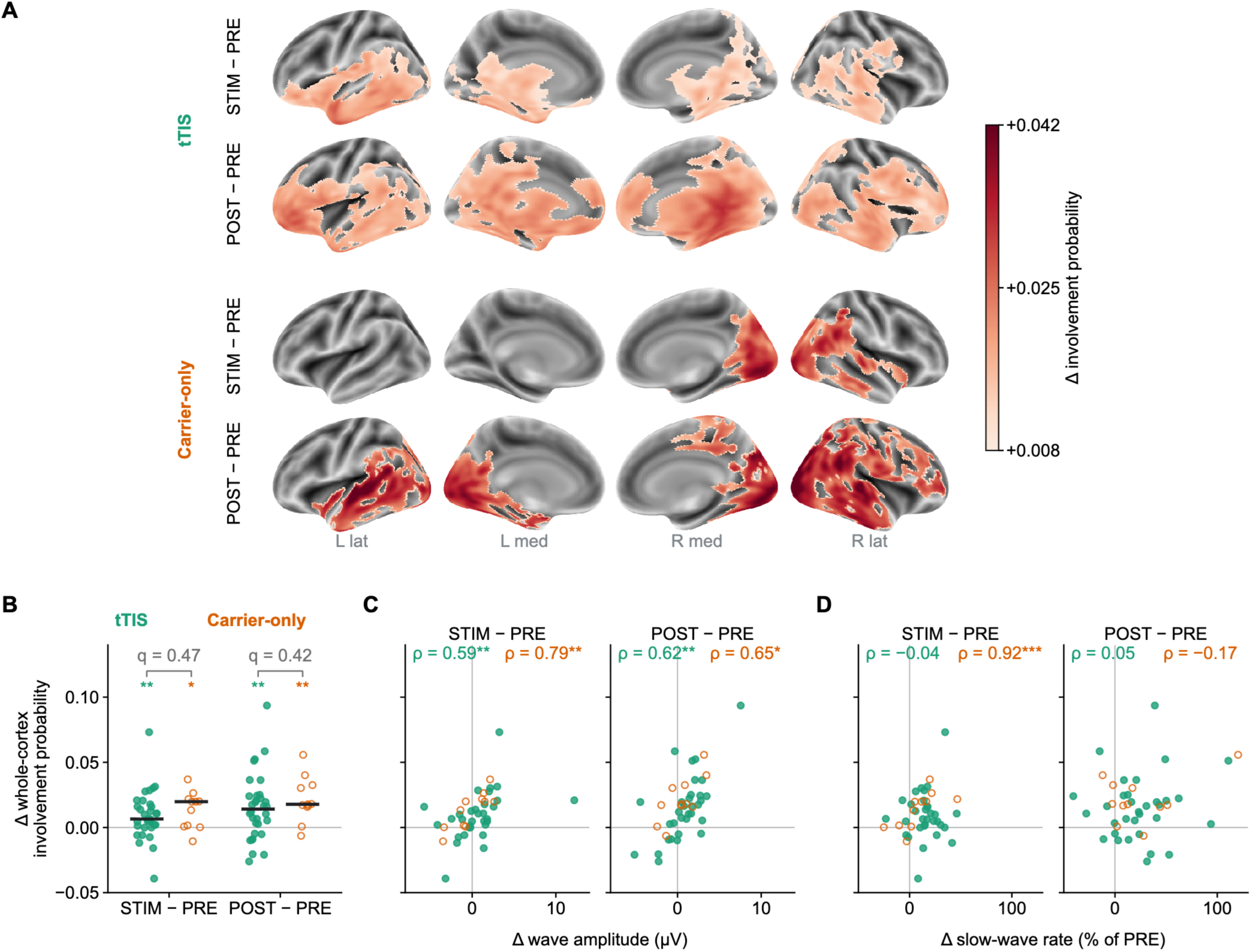
Source-space slow-wave involvement probability. (A) Change in the per-wave probability that a slow wave involves each vertex, STIM − PRE and POST − PRE, for tTIS and carrier-only on inflated fsaverage5. Maps show the group-mean change inside each group’s permutation-significant clusters (Wilcoxon signed-rank; cluster-forming p < 0.05, cluster mass, 5,000 sign flips, corrected p < 0.05); non-significant cortex is gray. Involvement probability rose in tTIS and carrier-only during and after stimulation. The groups did not differ. Origin probability did not change in either group or between them (Fig. S8; Table S4). (B) Change in involvement probability averaged over the whole cortex for each participant (points), with the median of each group (bar). The change was tested within each group (Wilcoxon signed-rank; asterisks) and between groups (Mann–Whitney; not significant, q above each pair). (C) The same change against the change in mean wave amplitude, taken as the negative-peak amplitude at each wave’s reference channel. (D) The same change against the change in slow-wave rate (merged waves per usable minute, percent of PRE). In C and D, ρ is the Spearman correlation within each group. Asterisks in B–D mark significance after Benjamini–Hochberg correction across the 14 tests of these panels (* q < 0.05, ** q < 0.01, *** q < 0.001). Filled points are tTIS and open points are carrier-only.

We next averaged involvement probability over the whole cortex for each participant, a measure that requires no cluster threshold. This average rose in tTIS (median change +0.007 during stimulation, q = 0.0067, and +0.014 after it, q = 0.0054) and in the carrier-only control (+0.020, q = 0.012, and +0.018, q = 0.0068), and the change did not differ between the groups (q = 0.47 and 0.42; Fig. 4B). Participants whose waves grew more in amplitude showed the larger rise in involvement, in tTIS (ρ = 0.59 for STIM/PRE and 0.62 for POST/PRE, both q ≤ 0.0017) and in the carrier-only control (ρ = 0.79, q = 0.0075, and 0.65, q = 0.0497; Fig. 4C). In tTIS the rise in involvement was unrelated to the change in slow-wave rate (ρ = −0.04 and 0.05, both q = 0.85), whereas in the carrier-only control the two were related during stimulation (ρ = 0.92, q < 0.001) but not after it (ρ = −0.17, q = 0.71; Fig. 4D). Each wave thus reached more cortex when the waves themselves were larger, in both groups, and in tTIS this change was separate from the gain in the number of waves. The wider involvement is therefore a property of the waves that both groups shared, and not a by-product of the density increase detected only in tTIS.

### Modeled Dose Tracks the Shift Toward N3 Sleep

The standardized montage delivered its strongest TI_normal_ to the left insula, which ranked first of 68 Desikan–Killiany regions by group mean, and fifth for the orientation-free TI_max_ (Fig. 5). Across tTIS participants, the inter-subject correlation of the whole-cortex field pattern was high for the two orientation-free magnitudes (mean Pearson r = 0.90 for both TI_max_ and HF_max_) but substantially lower for the surface-normal envelope (mean r = 0.48; Fig. 5C–F), reflecting its dependence on individual cortical folding.

**Fig. 5.**
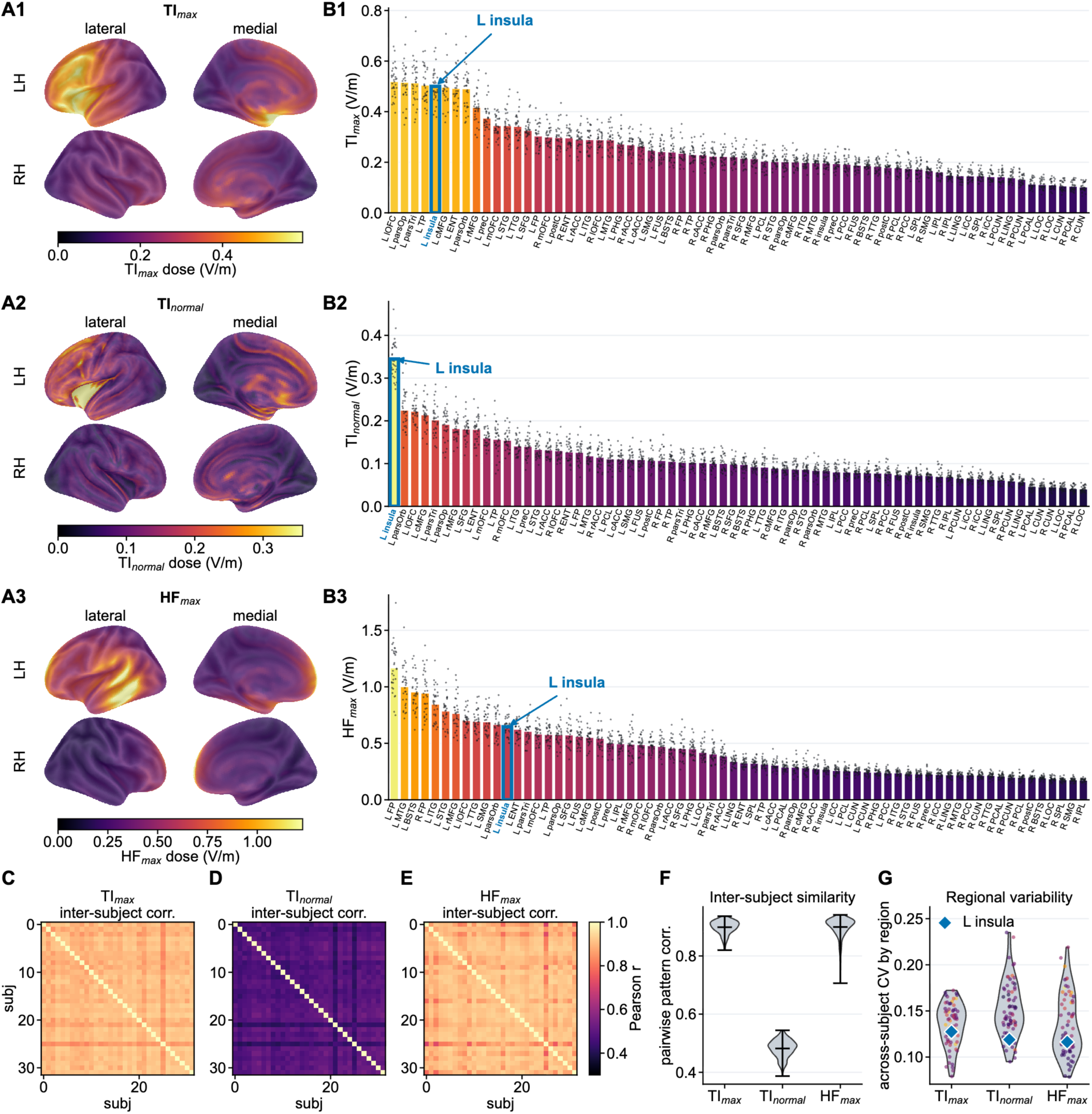
Modeled received dose and its inter-individual variability. (A1–A3) Group-mean modeled interference-field dose across the 32 tTIS participants on the inflated fsaverage5 surface (both hemispheres, lateral and medial) for the orientation-free envelope (TI_max_, A1), the surface-normal envelope (TI_normal_, A2), and the high-frequency carrier dose (HF_max_, A3). (B1–B3) Desikan–Killiany atlas regions ranked by group-mean dose for each metric, one dot per subject showing cohort variability, on the same per-metric scale; the left insula is highlighted. Region labels in B1–B3 give the hemisphere (L/R) and the Desikan–Killiany region. (C–E) Inter-subject field-pattern correlation matrices (Pearson r across cortical vertices between every pair of tTIS participants) for TI_max_ (C), TI_normal_ (D), and HF_max_ (E), on a shared scale. (F) Distribution of the pairwise pattern correlations per metric. (G) Across-subject coefficient of variation of field magnitude per Desikan–Killiany region (one point per region, colored by that region’s group-mean dose on the same color scale as A and B; the left insula is marked by a blue diamond).

The N3 share rose in both arms (N3 share in PRE, STIM and POST of 23.9, 26.0 and 28.3% in tTIS and 22.7, 24.7 and 26.6% in carrier-only). In the tTIS group, the magnitude of deepening, measured as the change in N3 share, followed the modeled field (Fig. 6A). Vertexwise, the change in N3 share from PRE during stimulation followed local TI_max_ in one left-hemisphere cluster that included perisylvian cortex and the insula (1,363 vertices; q = 0.011) and local HF_max_ in three frontal clusters (1,541 vertices; q = 0.011; Fig. 6C). Inside these clusters the mean field tracked the shift in tTIS (TI_max_ ρ = 0.69, 95% CI [0.44, 0.83]; HF_max_ ρ = 0.68, 95% CI [0.40, 0.84]) and not in the carrier-only control (ρ = −0.12 and −0.38). Inside these clusters, which were selected on the tTIS data, the dose relation also differed between the groups for both fields (descriptive p = 0.009 and 0.002).

**Fig. 6.**
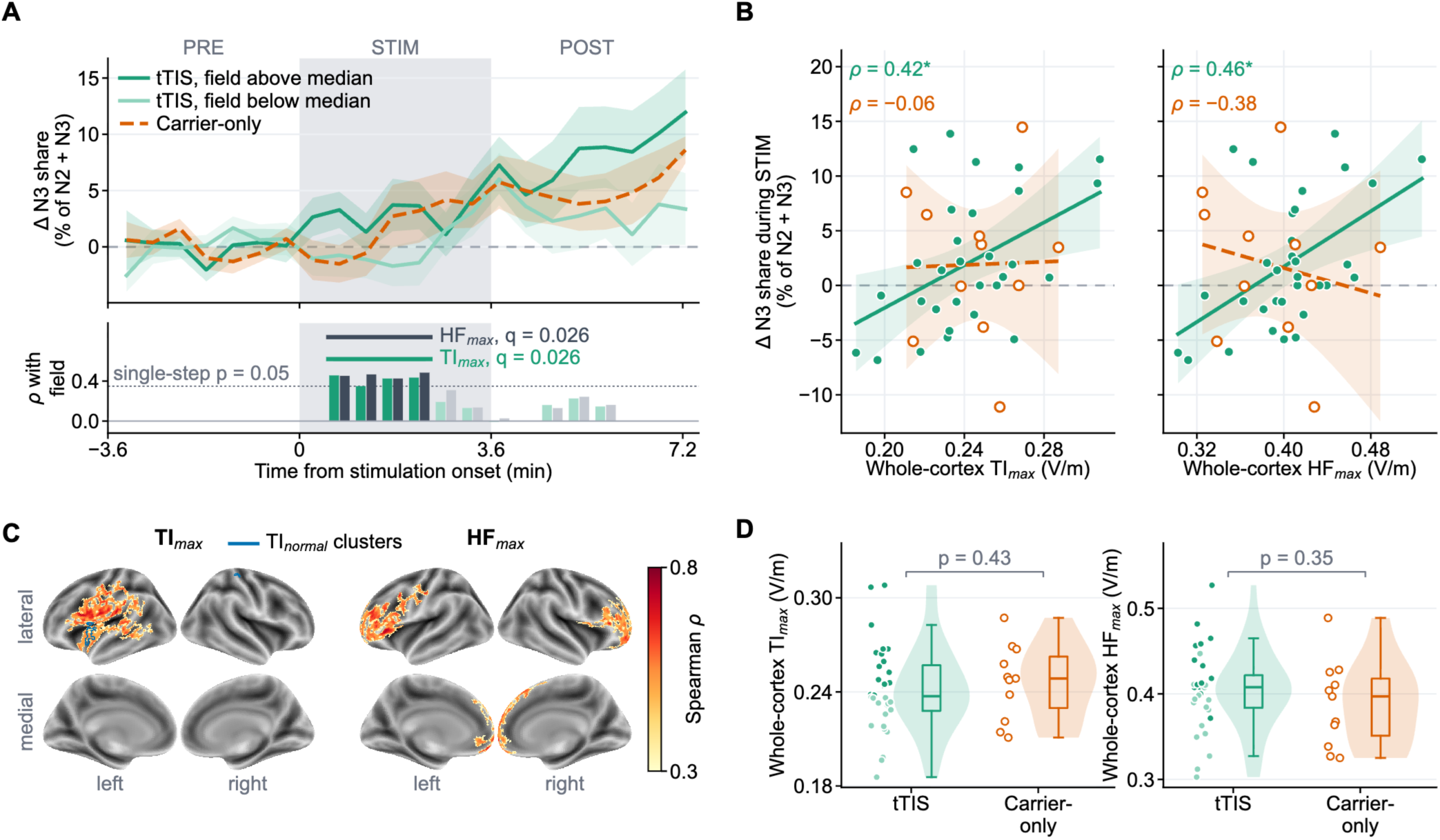
Modeled field and the shift toward N3 sleep during stimulation. (A) Change in N3 share from the PRE mean in 30-s steps aligned to stimulation onset (mean ± SEM), for tTIS participants above and below the median whole-cortex TI_max_ and for carrier-only; the split is for display and the gray band is the stimulation. Below, on the same time axis, the statistic that was tested: Spearman ρ across the 32 tTIS participants between the whole-cortex field (TI_max_, green; HF_max_, slate) and the change at each step that every participant reaches, with each field’s cluster of steps from a cluster-based permutation test over time (both 30 to 150 s, q = 0.026; full-color bars). Dotted, the ρ at which a single step reaches p = 0.05; pale steps are not significant on their own. Participant-level time courses are shown in Fig. S9. (B) Whole-cortex mean field, which involves no selection, against the change in N3 share averaged over the stimulation; tTIS filled, carrier-only open; least-squares fits with 95% confidence bands; Spearman ρ (* q < 0.05). (C) Spearman ρ, across the tTIS participants, of the local modeled field at each vertex with the change in N3 share during stimulation, shown inside significant clusters (cluster-forming p < 0.01): TI_max_, one left-hemisphere cluster including perisylvian cortex and the insula (cluster p = 0.002); HF_max_, three frontal clusters ( p ≤ 0.021); each map’s smallest cluster p gives q = 0.011. The color scale runs from ρ = 0.30 to 0.80. Blue outline, the three clusters of the separate vertexwise test of the surface-normal envelope TI_normal_ (56 vertices, p ≤ 0.045, q = 0.011), the largest in the left insula ( p = 0.002); 44 of the 56 vertices lie within the TI_max_ cluster (D) Whole-cortex mean field per group (violin; box, median and quartiles; points, participants; tTIS participants above the median TI_max_ dark green, the others light green, as split in A), with the between-group Mann–Whitney p. No envelope was delivered in carrier-only, so its TI_max_ is the modulation depth its two simulated carrier fields would have produced with a 1 Hz offset, a placebo, and its HF_max_ is the coherent field of its in-phase carriers.

With the whole-cortex mean field, which requires no cluster-based permutation test, the relation was weaker but present in tTIS (TI_max_ ρ = 0.42, 95% CI [0.04, 0.73], q = 0.029; HF_max_ ρ = 0.46, 95% CI [0.08, 0.75], q = 0.026; Fig. 6B) and not seen in carrier-only (ρ = −0.06 and −0.38), with a group difference for HF_max_ (q = 0.034) and not for TI_max_ (q = 0.22). The modeled field was higher in younger tTIS participants (ρ = −0.50 for TI_max_ and −0.48 for HF_max_), and after adjustment for age and sex, the relation remained for the cluster-mean field (ρ = 0.58 and 0.47, uncorrected p = 0.0008 and 0.0085) and weakened for the whole-cortex mean field (ρ = 0.28 and 0.33, uncorrected p = 0.13 and 0.072). In carrier-only no envelope was delivered, so its TI_max_ is a placebo that carries each participant’s anatomy on the same scale. Both groups received a similar modeled dose (Mann–Whitney p = 0.43 and 0.35; Fig. 6D). In time, the change in N3 share followed both TI_max_ and HF_max_ from 30 to 150 s after stimulation onset (q = 0.026 for each; Fig. 6A; participant-level time courses in Fig. S9), and after stimulation the correlations were not significant (ρ = 0.21 and 0.19). TI_normal_ formed three small clusters, the largest in the left insula (q = 0.011), and 44 of the 56 vertices lay inside the TI_max_ cluster (outline in Fig. 6C).

## Discussion

Non-invasive modulation of the deep cortical circuits that are a hot spot for sleep slow-wave generation has long been constrained by the physics of tES, which cannot reach a deep target without delivering even stronger fields to the overlying cortex. Here, tTIS targeting the left insula/ventromedial prefrontal cortex during NREM sleep increased detected slow waves during stimulation and that effect persisted afterward. In a received-dose analysis, tTIS participants who received a stronger modeled field shifted further toward N3 sleep during stimulation, a relation not seen in the carrier-only control.

Schaeffer et al. (2026) previously reported, using the same stimulation protocol in 28 participants (21 tTIS, 7 carrier-only), a subset of the 43 analyzed here, that tTIS increased spectral delta-band SWA during NREM sleep, an effect that outlasted stimulation ^16^. That result established spectral modulation and offered preliminary evidence of subjective benefit showing that, across the intervention period, the change in delta-band enhancement was positively associated with improved subjective ratings of sleep quality. The present study builds upon Schaeffer et al. (2026) and advances these lines into an analysis of discrete slow waves, their projection to source space, and a dose-response relationship.

### Sensor- and Source-Space Response

At the scalp, the slow-wave increase was broad and strongest for the density (number of waves per minute) during and after stimulation, spanning most of the EEG net and shallow and deep waves alike in the tTIS group, with no detected change in the carrier-only control (Fig. 3). Morphological changes were more limited, with modestly deeper negative peaks during stimulation. These changes were about 1% of each participant’s PRE value, against a density increase of 20– 30%, so tTIS changed how often slow waves occurred far more than their size or shape. The response was uneven across the night, as the gain was largest in the earlier stimulation episodes and waned across them (Fig. 3, Fig. S7). This waning sensitivity may reflect declining homeostatic sleep pressure, consistent with the synaptic homeostasis hypothesis linking overnight selective reductions in synaptic strength to declining slow-wave activity ^6^. Early in the night, when sleep pressure is high, cortical neurons are more synchronized and slow waves are larger and steeper ^20^, so a weak subthreshold input may more readily tip the network into additional slow waves. Averaging episodes across the night may therefore dilute an early response and should be considered in future tTIS sleep interventions. Due to our study design, episode order and elapsed time were nearly collinear, and so further insight regarding stimulation effects on homeostatic decline was limited (Fig. S7).

Source reconstruction extended our analysis to the cortex. Per wave, involvement rose during and after stimulation in both arms, with no difference between them (Fig. 4), while origin probability did not change (Fig. S8). This does not conflict with the scalp, where only tTIS showed a significant amplitude change. Those small amplitude changes did not differ between the arms (between-arm cluster p ≥ 0.18; Table S3), and involvement measures the extent of cortex active above its resting level at each wave’s negative peak rather than the wave’s amplitude, so in the carrier-only control it rose without a detectable change in wave amplitude (Fig. 4B,C). A rise present in both arms points, at least in part, to a factor common to them rather than to the envelope. One candidate is the natural deepening of sleep, as the N3 share rose from PRE to STIM and POST in both groups, and slow waves in deeper sleep are larger and more widespread ^20,21^. A second common factor is the kilohertz carriers, which both arms received, and without a night free of stimulation the two cannot be separated. Together, these maps suggest that tTIS added slow waves without changing where on the cortex they originate, and that the wider cortical involvement of each wave is not specific to the envelope. An unchanged origin fits the subthreshold regime of tTIS, unlike suprathreshold TMS pulses, which trigger slow waves at the stimulated site ^23^.

### Dose-Response Relationship

Sleep deepened across each stimulation episode in both groups, and in tTIS the participants who received a stronger modeled field shifted further toward N3 during stimulation (Fig. 6). The relation was not seen in the carrier-only control, whose in silico placebo TI_max_ carries the same anatomical variation without a delivered envelope. The relation held for the whole-cortex mean field and was strongest in a left-hemisphere cluster for TI_max_ and in frontal clusters for HF_max_. The two fields were too closely related across participants (ρ = 0.83 for their whole-cortex means) to attribute one and not the other. Age could confound this relation, because slow-wave sleep declines with age ^22^ and the modeled field depends on head anatomy that changes with age, such as the cerebrospinal fluid layer that widens with atrophy ^32,61^. Modeled field was indeed higher in younger participants (ρ = −0.50 for TI_max_ and −0.48 for HF_max_), and after adjustment for age and sex, the relation remained for the field inside the clusters (ρ = 0.58 and 0.47) but not for the whole-cortex mean field (ρ = 0.28 and 0.33), so age may account for part of the relation, and the part that remained rests on the cluster means. Because the field was modeled and not manipulated, these data show that field and deepening go together across people and not that one causes the other. If confirmed with the dose under experimental control, a field-dependent deepening of sleep would be of clinical interest, as deep NREM sleep declines with age and is disrupted in neurodegenerative and psychiatric disease ^4,22^, and it would make the individually modeled field a practical basis for personalizing stimulation.

The closest published tests relating an individually predicted tTIS dose to a response are hemodynamic. Violante et al. (2023) found that each participant’s modeled TI_max_ correlated weakly with their own hippocampal blood-oxygen-level-dependent (BOLD) change, whereas the HF_max_ did not ^31^. Here, the shift toward N3 followed envelope and carrier dose alike, in separate regions, so the present data cannot attribute it to the envelope rather than to the overall field. Modulation depth and carrier intensity therefore both remain candidate dose metrics, and an explicit dose-matching design, holding one fixed while varying the other, could test which waveform feature governs the response. To our knowledge, this is the first report relating an individually modeled tES dose directly to the deepening of sleep.

While slow waves are known to be distributed traveling events ^19,21,29^, the insula remains a candidate intervention target for sleep modulation via tTIS. The montage delivered its strongest surface-normal envelope TI_normal_ to the left insula (Fig. 5), yet TI_normal_ formed only a small cluster there with the shift toward N3 (Fig. 6C). The normal component nonetheless merits modeling, as cellular and human tES evidence indicates that the field component along the somatodendritic axis, approximated by the surface normal, governs somatic polarization ^62–64^. Similarly in mice, the orientation of matched envelope amplitude interference fields relative to hippocampal axons determined whether stimulation evoked seizure-like activity ^65^. This orientation dependence is notable because, unlike the tDCS or tACS field, which at each location acts along a fixed axis, the tTIS field rotates over the beat cycle ^66^. To our knowledge, how neurons respond to the direction of such a field relative to their morphology has been examined only in silico and in vivo ^65,66^, and in vitro preparations, in which the field direction can be set relative to the cell, could test it under controlled conditions. TI_normal_, whose cortical projection follows each person’s gyral geometry, is also the most inter-individually variable field metric (Fig. 5C–G), which group-average or magnitude-only models obscure ^30,32,67^.

A dose-response association that is present in tTIS and absent in the carrier-only control at a comparable intensities, as seen here, would be consistent with the beat acting as the gate while carrier dose scales the response. Such a beat dependence would resemble prior work in which kilohertz carriers drove envelope-locked firing in mice only when offset to produce a beat and were inert without one ^11^. Similarly, in in silico models of cortical neurons, tTIS evoked spikes locked to the beat, whereas fields dominated by one carrier, with little modulation, were less effective at evoking spikes ^66^. However, both these studies used suprathreshold fields that drove spiking. Recent work extends this beat dependence below spiking threshold. In mouse cortical slices, subthreshold kilohertz carriers alone depolarized neurons without temporal structure, whereas a 1 Hz envelope added membrane-potential oscillations, although the field strength in these experiments was not reported ^68^. This is consistent with the pattern in our arms, although the between-arm density difference during stimulation (cluster p = 0.030) did not survive correction across maps. Carrier-only stimulation produced no detected change in slow-wave density and no detected dose association, whereas in tTIS the same carriers with a 1 Hz beat increased slow-wave density. A depolarization without temporal structure may thus be insufficient to change how often slow waves occur, whereas a periodic component near the slow-wave frequency may be, which leaves open how the beat acts on the sleeping cortex.

### Open Questions and Future Directions

The mechanism underlying the tTIS-induced increase in slow waves remains unresolved. Nonlinear neuronal responses and interactions with ongoing network activity have been proposed to contribute to tTIS effects ^10,69–72^, but much of this evidence comes from suprathreshold fields, and how kilohertz fields act on neurons in the subthreshold regime remains contested. One question is whether stimulation simply raises slow-wave propensity via membrane potential biasing or larger network entrainment to the exogenous signal. The increase in wave density across shallow and deep waves alike, with only modest morphology change, points to the former. Such a biasing effect is plausible at weak field strengths, as in cortical slices generating the slow oscillation, applied fields within the range of endogenous field strengths, which change the membrane potential of single neurons only slightly, enhanced and entrained network activity ^73^. Evidence from tACS modeling and primate tTIS recordings suggests that external stimulation can either disrupt ongoing synchrony or promote phase locking, depending on stimulation parameters and state of endogenous activity ^74,75^. In tACS models, local fields below 0.3 V/m can desynchronize neuronal firing from ongoing oscillations, whereas fields above 0.3 V/m can promote entrainment to the applied stimulus, and entrainment requires the least field strength when the stimulation frequency matches the endogenous rhythm ^74^. Primate tTIS recordings likewise show bidirectional changes in spike timing without changes in firing rate. Phase locking mostly decreased in neurons strongly locked at baseline and increased in weakly locked ones, with slightly more entrainment at higher amplitude ^75^.

Such a dependence on field strength suggests one possible interpretation of the dose relation observed here. In primates, weak tACS competes with ongoing oscillations for control of spike timing and reduces the entrainment of neurons that were locked to them, whereas stronger stimulation imposes its own timing ^76^. By analogy, a weak envelope field might interfere with the transition into deeper sleep and a stronger one might facilitate it, whereas carriers without an envelope would have neither effect. The shift toward N3 during stimulation increased with the modeled field in tTIS and not in the carrier-only control (Fig. 6), which is consistent with this interpretation. Future studies could vary envelope frequency around each participant’s endogenous rhythm and stimulation amplitude, while accounting for the resulting local field strength, to test for an Arnold tongue ^74^. Combined with direct phase-locking measurements, this could help distinguish entrainment from a broader increase in slow-wave propensity. In such cases, a design aspect worth considering is the length of the stimulation windows, since detecting entrainment with phase-coherence measures depends critically on epoch length ^77^.

Treating the individually modeled field as the dose received by the brain assumes the predicted fields by in silico models track delivered fields, an assumption partially supported by initial intracranial recordings ^78,79^. Prior measurements in two primates showed correspondence in spatial field distributions (Spearman ρ ≈ 0.7), but differed with respect to field intensity (recorded-versus-predicted regression slopes s = 0.50 and 1.12) ^79^. Human stereo-EEG confirmed that tTIS engages deep targets and dissociates envelope-from carrier-driven effects ^78^. However, a formal validation of the accuracy of models against direct measurements in a larger human study, similar to those conducted for earlier tES modalities in humans, is still warranted ^80,81^. Such investigation would be useful to establish accurate and reliable dose-response relationships from computational models ^53^.

## Conclusion

This study shows that 1 Hz tTIS delivered during NREM sleep increased the density of discrete slow waves during stimulation, an increase that persisted after stimulation ended. Sleep depth also followed the received dose, in that the stronger the modeled field a tTIS participant received, the further their sleep shifted toward N3 during stimulation, a relation not seen in the carrier-only control. Together, these results extend previous tTIS sleep research from spectral power to discrete slow-wave events, their cortical distribution, and the field each brain received. Beyond these findings, tTIS adds a much-needed tool for sleep research. It can be delivered below perception threshold during sleep ^16^, it permits concurrent EEG with suitable hardware filtering ^15,16,82^, and it can reach deep targets that conventional tES cannot engage without stronger fields to the overlying cortex ^11,31^. It therefore offers a practical way to test how deep regions contribute to slow waves and to the restorative functions of sleep. If the dose dependence is confirmed, stimulation could be tailored to the individual in the many conditions in which deep sleep is diminished, and the compact hardware of tTIS suggests a path toward home use, as already considered for other forms of tES ^83^.

## Acknowledgments

We thank the participants who volunteered for this study for their time and contribution. We thank the staff at the University of Wisconsin–Madison who contributed to participant recruitment (Tricia Denman, Emily Peterson, and Kate Riordan), data collection (Everett Carstens, Claire Doucet, Paul Ladefoged, Poorang Nori, Richard Smith, and Ana Maria Vascan), and data processing (Cameron Brace and Dillon Scott), and the staff at the Waisman Center Brain Imaging Laboratory and the Center for Healthy Minds, where the MRI data were collected. The TIBS-R investigational devices used in this project were provided by TI Solutions AG, Switzerland, as part of its Early Adopter Program (www.temporalinterference.com). The content of the information does not necessarily reflect the position or the policy of the Government, and no official endorsement should be inferred.

## Data Availability

Summarized source data for the main data figures and table are available at https://github.com/idossha/tTIS_sw_reconstruction. Source data for the Supplementary Information and unprocessed data are available from the corresponding author upon reasonable request. TI-Toolbox is available at https://github.com/idossha/TI-Toolbox.

## Declaration of generative AI and AI-assisted technologies in the manuscript preparation process

During the preparation of this work, the authors used Claude Opus 5.5 (Anthropic) in order to assist with text editing, restructuring, and clarity improvements. No scientific content was generated by AI. After using this tool/service, the authors reviewed and edited the content as needed and take full responsibility for the content of the published article.

## Author Contributions

Conceptualization – I.H., G.T.; Data Curation – I.H., E.L.S., Z.F.; Formal Analysis – I.H., E.L.S.; Funding Acquisition – G.T., S.G.J.; Investigation – E.L.S., I.H., B.M.; Methodology – I.H., E.L.S., Z.F., G.T., S.G.J., M.B.; Project Administration – S.G.J., E.L.S., B.M., M.B.; Software – I.H., E.L.S., Z.F.; Visualization – I.H.; Writing (Original Draft) – I.H., G.T.; Writing (Review & Editing) – all authors.

## References

1. Tononi, G. & Cirelli, C. Sleep and the price of plasticity: from synaptic and cellular homeostasis to memory consolidation and integration. Neuron 81, 12–34 (2014).

2. Krause, A. J. et al. The sleep-deprived human brain. Nature Reviews Neuroscience 18, 404–418 (2017).

3. Fehér, K. D. et al. Shaping the slow waves of sleep: A systematic and integrative review of sleep slow wave modulation in humans using non-invasive brain stimulation. Sleep Medicine Reviews 58, 101438 (2021).

4. Wulff, K., Gatti, S., Wettstein, J. G. & Foster, R. G. Sleep and circadian rhythm disruption in psychiatric and neurodegenerative disease. Nature Reviews Neuroscience 11, 589–599 (2010).

5. Rasch, B. & Born, J. About sleep’s role in memory. Physiological Reviews 93, 681–766 (2013).

6. Tononi, G. & Cirelli, C. Sleep function and synaptic homeostasis. Sleep Medicine Reviews 10, 49–62 (2006).

7. Bellesi, M., Riedner, B. A., Garcia-Molina, G. N., Cirelli, C. & Tononi, G. Enhancement of sleep slow waves: underlying mechanisms and practical consequences. Frontiers in Systems Neuroscience 8, 208 (2014).

8. Polanía, R., Nitsche, M. A. & Ruff, C. C. Studying and modifying brain function with non-invasive brain stimulation. Nature Neuroscience 21, 174–187 (2018).

9. Helfrich, R. F. et al. Entrainment of brain oscillations by transcranial alternating current stimulation. Current Biology 24, 333–339 (2014).

10. Esmaeilpour, Z., Kronberg, G., Reato, D., Parra, L. C. & Bikson, M. Temporal interference stimulation targets deep brain regions by modulating neural oscillations. Brain Stimulation 14, 55–65 (2021).

11. Grossman, N. et al. Noninvasive deep brain stimulation via temporally interfering electric fields. Cell 169, 1029–1041.e16 (2017).

12. Albantakis, L. & Tononi, G. Precision neuromodulation in psychiatry: focus on temporal interference stimulation. American Journal of Psychiatry 183, 721–733 (2026).

13. Demchenko, I. et al. Human applications of transcranial temporal interference stimulation: a systematic review. Brain Stimulation 18, 2054–2066 (2025).

14. Wang, F., Jiang, M., Deng, D. & Tu, Y. Temporal interference stimulation for human brain: opportunities and challenges. The Innovation 6, 100803 (2025).

15. von Conta, J., Kasten, F. H., Ćurčić-Blake, B., Schellhorn, K. & Herrmann, C. S. Characterizing low-frequency artifacts during transcranial temporal interference stimulation (tTIS). Neuroimage: Reports 2, 100113 (2022).

16. Schaeffer, E. L. et al. Enhancement of sleep slow wave activity using transcranial electrical stimulation with temporal interference: an interim analysis of the STRENGTHEN study. Communications Medicine 6, 549 (2026).

17. Thiele, C., Tamm, C., Ruhnau, P. & Zaehle, T. Perceptibility and pain thresholds in low- and high-frequency alternating current stimulation: implications for tACS and tTIS. Journal of Cognitive Enhancement 9, 79–91 (2025).

18. Steriade, M., Nunez, A. & Amzica, F. A novel slow (¡ 1 Hz) oscillation of neocortical neurons in vivo: depolarizing and hyperpolarizing components. Journal of Neuroscience 13, 3252–3265 (1993).

19. Massimini, M., Huber, R., Ferrarelli, F., Hill, S. & Tononi, G. The sleep slow oscillation as a traveling wave. Journal of Neuroscience 24, 6862–6870 (2004).

20. Riedner, B. A. et al. Sleep homeostasis and cortical synchronization III: a high-density EEG study of sleep slow waves in humans. Sleep 30, 1643–1657 (2007).

21. Nir, Y. et al. Regional slow waves and spindles in human sleep. Neuron 70, 153–169 (2011).

22. Carrier, J. et al. Sleep slow wave changes during the middle years of life. European Journal of Neuroscience 33, 758–766 (2011).

23. Massimini, M. et al. Triggering sleep slow waves by transcranial magnetic stimulation. Proceedings of the National Academy of Sciences of the United States of America 104, 8496–8501 (2007).

24. Ngo, H.-V. V., Martinetz, T., Born, J. & Molle, M. Auditory closed-loop stimulation of the sleep slow oscillation enhances memory. Neuron 78, 545–553 (2013).

25. Marshall, L., Helgadottir, H., Molle, M. & Born, J. Boosting slow oscillations during sleep potentiates memory. Nature 444, 610– 613 (2006).

26. Hathaway, E. et al. Transcranial electrical stimulation targeting limbic cortex increases the duration of human deep sleep. Sleep Medicine 81, 350–357 (2021).

27. Lafon, B. et al. Low frequency transcranial electrical stimulation does not entrain sleep rhythms measured by human intracranial recordings. Nature Communications 8, 1199 (2017).

28. Bruno, S. et al. Thalamic transcranial electrical stimulation with temporal interference enhances sleep spindle activity during a daytime nap. SLEEP Advances 7, zpag069 (2026).

29. Murphy, M. et al. Source modeling sleep slow waves. Proceedings of the National Academy of Sciences of the United States of America 106, 1608–1613 (2009).

30. Kasten, F. H., Duecker, K., Maack, M. C., Meiser, A. & Herrmann, C. S. Integrating electric field modeling and neuroimaging to explain inter-individual variability of tACS effects. Nature Communications 10, 5427 (2019).

31. Violante, I. R. et al. Non-invasive temporal interference electrical stimulation of the human hippocampus. Nature Neuroscience 26, 1994–2004 (2023).

32. Haber, I., Jackson, A., Thielscher, A., Hai, A. & Tononi, G. TI-Toolbox: An open-source software for temporal interference stimulation research. Brain Stimulation 19, 103016 (2026).

33. University of Wisconsin, Madison. Hybrid Interventions to Optimize Neural Plasticity and Enhance Well-being: Mental Exercise, and Transcranial Electrical Stimulation With Temporal Interference: The STRENGTHEN Study - Phase 1 and 2. (2024).

34. Franke, M., Bhadra, N., Bhadra, N. & Kilgore, K. Direct current contamination of kilohertz frequency alternating current waveforms. Journal of Neuroscience Methods 232, 74–83 (2014).

35. Troester, M. M., Quan, S. F., Berry, R. B. et al. The AASM Manual for the Scoring of Sleep and Associated Events: Rules, Terminology and Technical Specifications, Version 3. (American Academy of Sleep Medicine, 2023).

36. Delorme, A. & Makeig, S. EEGLAB: an open source toolbox for analysis of single-trial EEG dynamics including independent component analysis. Journal of Neuroscience Methods 134, 9–21 (2004).

37. Palmer, J. A., Kreutz-Delgado, K. & Makeig, S. AMICA: An Adaptive Mixture of Independent Component Analyzers with Shared Components. (2012).

38. Noury, N., Hipp, J. F. & Siegel, M. Physiological processes non-linearly affect electrophysiological recordings during transcranial electric stimulation. NeuroImage 140, 99–109 (2016).

39. Perrin, F., Pernier, J., Bertrand, O. & Echallier, J. F. Spherical splines for scalp potential and current density mapping. Electroencephalography and Clinical Neurophysiology 72, 184–187 (1989).

40. Gramfort, A. et al. MEG and EEG data analysis with MNE-Python. Frontiers in Neuroscience 7, 267 (2013).

41. Vallat, R. & Walker, M. P. An open-source, high-performance tool for automated sleep staging. eLife 10, e70092 (2021).

42. Mensen, A., Riedner, B. & Tononi, G. Optimizing detection and analysis of slow waves in sleep EEG. Journal of Neuroscience Methods 274, 1–12 (2016).

43. Puonti, O. et al. Accurate and robust whole-head segmentation from magnetic resonance images for individualized head modeling. NeuroImage 219, 117044 (2020).

44. Fischl, B. FreeSurfer. NeuroImage 62, 774–781 (2012).

45. Ledoit, O. & Wolf, M. A well-conditioned estimator for large-dimensional covariance matrices. Journal of Multivariate Analysis 88, 365–411 (2004).

46. Engemann, D. A. & Gramfort, A. Automated model selection in covariance estimation and spatial whitening of MEG and EEG signals. NeuroImage 108, 328–342 (2015).

47. Pascual-Marqui, R. D. Standardized low-resolution brain electromagnetic tomography (sLORETA): technical details. Methods and Findings in Experimental and Clinical Pharmacology 24, 5–12 (2002).

48. Haber, I., Jackson, A., Thielscher, A., Hai, A. & Tononi, G. TI-Toolbox vv3.1.0. 10.5281/zenodo.23268477 (2026).

49. Cassarà, A. M. et al. Recommendations for the Safe Application of Temporal Interference Stimulation in the Human Brain Part I: Principles of Electrical Neuromodulation and Adverse Effects. Bioelectromagnetics 46, e22542 (2025).

50. Haber, I., Tononi, G. & Albantakis, L. Same currents, different field: Δf=0 carrier-only stimulation is not temporal interference without the beat. arXiv preprint arXiv:2611.14839 (2026).

51. Saturnino, G. B. et al. in Brain and Human Body Modeling: Computational Human Modeling at EMBC 2018 3–25 (Springer, 2019).

52. Van Hoornweder, S. et al. Outcome measures for electric field modeling in tES and TMS: A systematic review and large-scale modeling study. NeuroImage 281, 120379 (2023).

53. Soleimani, G. et al. Dose-response relationships in transcranial brain stimulation: Physics, physiology and mechanism. Brain Stimulation 19, 103067 (2026).

54. Maris, E. & Oostenveld, R. Nonparametric statistical testing of EEG- and MEG-data. Journal of Neuroscience Methods 164, 177– 190 (2007).

55. Benjamini, Y. & Hochberg, Y. Controlling the False Discovery Rate: A Practical and Powerful Approach to Multiple Testing. Journal of the Royal Statistical Society Series B: Statistical Methodology 57, 289–300 (1995).

56. Kerby, D. S. The simple difference formula: an approach to teaching nonparametric correlation. Comprehensive Psychology 3, 11.IT.3.1 (2014).

57. Wendt, H. W. Dealing with a common problem in social science: a simplified rank-biserial coefficient of correlation based on the U statistic. European Journal of Social Psychology 2, 463–465 (1972).

58. Saravanan, V., Berman, G. J. & Sober, S. J. Application of the hierarchical bootstrap to multi-level data in neuroscience. *Neurons, Behavior*, Data analysis, and Theory 3, 1–25 (2020).

59. Baetens, K., Van Hoornweder, S., Berger, T. A. & Wischnewski, M. ACES: Automated Correlation of Electric field strength and Stimulation effects for non-invasive brain stimulation. Brain Stimulation 17, 473–475 (2024).

60. Desikan, R. S. et al. An automated labeling system for subdividing the human cerebral cortex on MRI scans into gyral based regions of interest. NeuroImage 31, 968–980 (2006).

61. Indahlastari, A. et al. Modeling transcranial electrical stimulation in the aging brain. Brain stimulation 13, 664–674 (2020).

62. Radman, T., Ramos, R. L., Brumberg, J. C. & Bikson, M. Role of cortical cell type and morphology in subthreshold and suprathreshold uniform electric field stimulation in vitro. Brain Stimulation 2, 215–228 (2009).

63. Rahman, A. et al. Cellular effects of acute direct current stimulation: somatic and synaptic terminal effects. The Journal of Physiology 591, 2563–2578 (2013).

64. Laakso, I., Mikkonen, M., Koyama, S., Hirata, A. & Tanaka, S. Can electric fields explain inter-individual variability in transcranial direct current stimulation of the motor cortex? Scientific Reports 9, 626 (2019).

65. Missey, F. et al. Orientation of Temporal Interference for Non-invasive Deep Brain Stimulation in Epilepsy. Frontiers in Neuroscience 15, 633988 (2021).

66. Wang, B., Aberra, A. S., Grill, W. M. & Peterchev, A. V. Responses of model cortical neurons to temporal interference stimulation and related transcranial alternating current stimulation modalities. Journal of Neural Engineering 19, 066047 (2022).

67. Laakso, I., Tanaka, S., Koyama, S., De Santis, V. & Hirata, A. Inter-subject variability in electric fields of motor cortical tDCS. Brain Stimulation 8, 906–913 (2015).

68. Yoshimoto, R., et al. Kilohertz electrical stimulation drives widespread neuronal activation and temporal interference imposes focal rhythmic modulation. bioRxiv (2026).

69. Opančar, A. et al. The same biophysical mechanism is involved in both temporal interference and direct kHz stimulation of peripheral nerves. Nature Communications 16, 9006 (2025).

70. Luff, C. E. et al. The neuron mixer and its impact on human brain dynamics. Cell Reports 43, 114274 (2024).

71. Mirzakhalili, E., Barra, B., Capogrosso, M. & Lempka, S. F. Biophysics of temporal interference stimulation. Cell Systems 11, 557–572.e5 (2020).

72. Wang, Y. et al. Advances in transcranial alternating current stimulation and temporal interference stimulation for neuropsychiatric disorders: mechanisms, technical developments, and preclinical evidence. Translational Psychiatry (2026).

73. Fröhlich, F. & McCormick, D. A. Endogenous Electric Fields May Guide Neocortical Network Activity. Neuron 67, 129–143 (2010).

74. Zhao, Z., Shirinpour, S., Tran, H., Wischnewski, M. & Opitz, A. Intensity- and frequency-specific effects of transcranial alternating current stimulation are explained by network dynamics. Journal of Neural Engineering 21, 026024 (2024).

75. Vieira, P. G., Krause, M. R. & Pack, C. C. Temporal interference stimulation disrupts spike timing in the primate brain. Nature Communications 15, 4558 (2024).

76. Krause, M. R., Vieira, P. G., Thivierge, J. & Pack, C. C. Brain stimulation competes with ongoing oscillations for control of spike timing in the primate brain. PLoS Biology 20, e3001650 (2022).

77. Mc Laughlin, M., Khatoun, A. & Asamoah, B. Detection of tACS entrainment critically depends on epoch length. Frontiers in Cellular Neuroscience 16, 806556 (2022).

78. Missey, F. et al. Non-invasive temporal interference stimulation of the hippocampus suppresses epileptic biomarkers in patients with epilepsy: biophysical differences between kilohertz and amplitude modulated stimulation. Brain Stimulation 19, 102981 (2026).

79. Liu, R. et al. Temporal interference stimulation targets deep primate brain. NeuroImage 291, 120581 (2024).

80. Huang, Y. et al. Measurements and models of electric fields in the in vivo human brain during transcranial electric stimulation. eLife 6, e18834 (2017).

81. Opitz, A. et al. Spatiotemporal structure of intracranial electric fields induced by transcranial electric stimulation in humans and nonhuman primates. Scientific Reports 6, 31236 (2016).

82. Rhodes, E., et al. Artefact-free EEG during temporal interference stimulation reveals phase-specific modulation of the human visual cortex. bioRxiv (2026).

83. Antal, A. et al. Low intensity transcranial electric stimulation: Safety, ethical, legal regulatory and application guidelines. Clinical Neurophysiology 128, 1774–1809 (2017).

