## Supplementary material for "Transcranial temporal interference stimulation increases slow waves during human NREM sleep": SI

### Contents


- Stimulation delivery and recording quality
- Sleep architecture of the stimulation night
- Assessment of residual 1 Hz stimulation artifact
- Participant-scaled slow-wave detection
- Cluster-based permutation statistics
- Sensor-space findings and within-night changes
  - Detectable effect sizes
  - Time-of-night dependence of wave morphology
  - Within-night decline: protocol order versus cleaned-recording time
- Source-space findings
  - Source origin and involvement probability maps
- Participant time courses of the shift toward N3 sleep
- Leave-one-subject-out sensitivity

### Stimulation delivery and recording quality

We summarized TI Brain Stimulator (TIBS-R v3.0) stimulation logs of delivered intensities through the hardware channels (tTIS: 1 Hz offset, 15,000/15,001 Hz; carrier-only: 0 Hz offset, 15,000/15,000 Hz), calculating plateau current and stimulation impedance per channel over each participant's logged protocols. Current clustered near the 5 mA target, with comparable, low impedance in both arms (Fig. S1, Table S1). One tTIS participant was titrated to 4 mA; another had asymmetric delivery (~6.3/4.0 mA). Importantly, field modeling used participant-specific channel-current inputs. Current and impedance were available for 42 of 43 participants. The participant without usable logs was excluded from this summary; field modeling assumed 5.15 mA per channel for that participant.

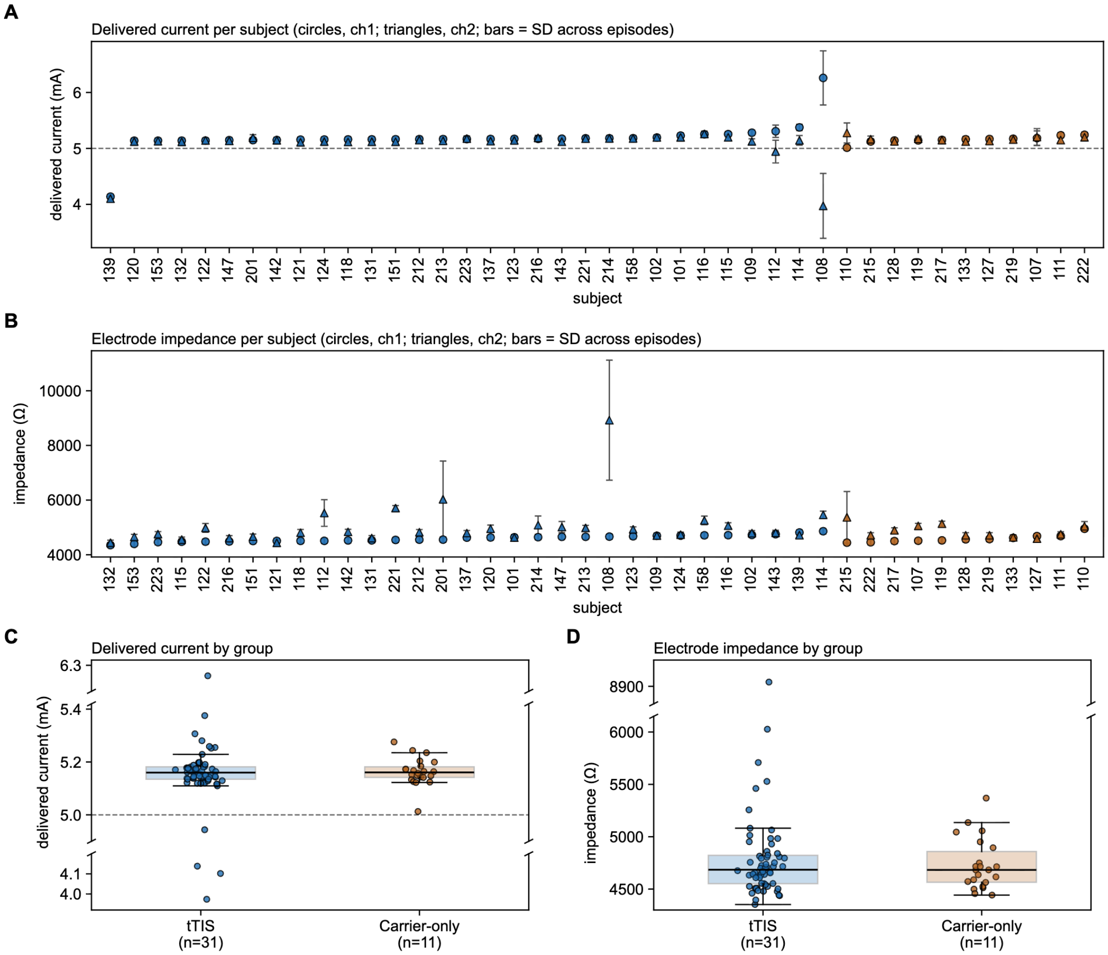


**Fig. S1.** *Delivered current and stimulation impedance per participant. (A) Participant-mean current by channel (circles, channel 1; triangles, channel 2; dashed line, 5 mA target). Error bars show episode SDs, with a minimum visible length; true SDs are in the per-subject data. (B) Participant-mean impedance by channel, with the same error-bar convention. (C,D) Current and impedance by group (boxes, median and interquartile range; points, participant×channel means); their y-axes are broken (diagonal marks) so the bulk of each distribution is resolved, and outlying participant×channel means are shown beyond the breaks. Of 32 tTIS and 11 carrier-only participants, measured current and impedance were available for 31 and 11, respectively.*

**Table S1.** *Delivered current and stimulation impedance by group (mean ± SD across participant-by-channel means).*

| Group | Subjects # | Protocols | Delivered current (mA) | Impedance (kΩ) |
|---|---|---|---|---|
| tTIS | 31 / 32 | 309 | 5.13 ± 0.28 | 4.82 ± 0.61 |
| Carrier-only | 11 / 11 | 105 | 5.16 ± 0.05 | 4.73 ± 0.24 |
| All | 42 / 43 | 414 | 5.14 ± 0.25 | 4.80 ± 0.54 |

### Sleep architecture of the stimulation night

Sleep architecture of the stimulation night was similar in the two arms (Table S2). Total sleep time, sleep efficiency and the minutes and share of each sleep stage did not differ. The sleep period was somewhat longer in tTIS (uncorrected p = 0.02), a difference that did not survive correction across the table (q = 0.23).

**Table S2.** *Sleep architecture of the stimulation night by arm (mean ± SD).*

| Measure | tTIS (n = 31) | Carrier-only (n = 11) | p |
|---|---|---|---|
| Sleep period time (min) | 442 ± 49 | 409 ± 27 | 0.02 |
| Total sleep time (min) | 385 ± 60 | 367 ± 61 | 0.31 |
| Wake after sleep onset (min) | 57.3 ± 62.4 | 42.0 ± 52.6 | 0.19 |
| Sleep efficiency (%) | 87.5 ± 12.7 | 89.7 ± 12.9 | 0.23 |
| N1 (min) | 16.9 ± 15.7 | 12.2 ± 12.2 | 0.32 |
| N2 (min) | 237.8 ± 45.8 | 236.5 ± 36.9 | 0.77 |
| N3 (min) | 47.7 ± 27.1 | 42.9 ± 26.2 | 0.74 |
| REM (min) | 82.1 ± 23.8 | 75.2 ± 30.2 | 0.94 |
| N1 (% of sleep) | 4.6 ± 4.3 | 3.9 ± 4.9 | 0.49 |
| N2 (% of sleep) | 61.8 ± 5.5 | 65.1 ± 8.3 | 0.35 |
| N3 (% of sleep) | 12.6 ± 6.9 | 11.2 ± 6.5 | 0.61 |
| REM (% of sleep) | 21.0 ± 5.8 | 19.8 ± 6.0 | 0.55 |
| Analyzed stimulation episodes | 9.7 ± 1.6 | 9.2 ± 2.2 | 0.93 |

*Note. Sleep was scored in 30-s epochs. Sleep period time runs from sleep onset (the first epoch scored N1 or N2) to the end of the last epoch scored as sleep. Wake after sleep onset is the part of the sleep period not scored as sleep, and includes unscored stretches, so it is an upper bound. Sleep efficiency is total sleep time divided by sleep period time. Stage percentages are of total sleep time. The recordings hold no lights-off marker, so time in bed and sleep onset latency are not reported. One tTIS participant is left out because sleep scoring of that recording ends after the first 2 h. Arms were compared with Mann–Whitney U tests (two-sided); p is uncorrected, and no measure differed after Benjamini–Hochberg correction across the 13 rows (all q ≥ 0.23).*

### Assessment of residual 1 Hz stimulation artifact

The two kHz carriers in tTIS produce a 1 Hz envelope. A residual envelope, DC offset [1], or nonlinear intermodulation product [2] could introduce slow-wave-band artifact and inflate detected wave counts during stimulation.

Several features reduce this concern. A 7th-order 100 Hz low-pass filter before the EEG amplifier attenuated carriers; a 4th-order 250 Hz high-pass filter on the stimulator output attenuated low-frequency components generated upstream (Methods). Wave morphology changed only modestly, with slope and frequency unchanged in tTIS during stimulation (Table S3), whereas an injected low-frequency waveform could alter detected wave shape. The increase also persisted after current delivery ended, supporting a physiological component without excluding concurrent STIM artifact.

We also tested the recorded data. For each participant, per-channel Welch spectra were estimated in every PRE, STIM, and POST interval using 60 s windows (0.0167 Hz bins), averaged across protocols and channels, then across participants. In tTIS, slow-wave-band (0.5–1.5 Hz) power exceeded PRE during STIM (Wilcoxon $p = 3.1 \times 10^{-6}$) and POST ($p = 2.0 \times 10^{-4}$); in the carrier-only arm it did so after (p = 0.024) but not during stimulation (p = 0.24). A stable 1 Hz artifact would appear as a narrow line at the 1 Hz bin. Instead, the increase was broadband. The three bins centered at 1 Hz held 6.7% of the band increase across all 183 channels and 7.6% in the 23 channels surrounding the stimulation electrodes (Fig. S2). In the no-beat carrier-only arm (15,000/15,000 Hz; n = 11), the same three bins held 7.9% of the band increase, but its STIM − PRE difference was narrower than in tTIS (full width at half maximum 0.18 versus 0.35 Hz).

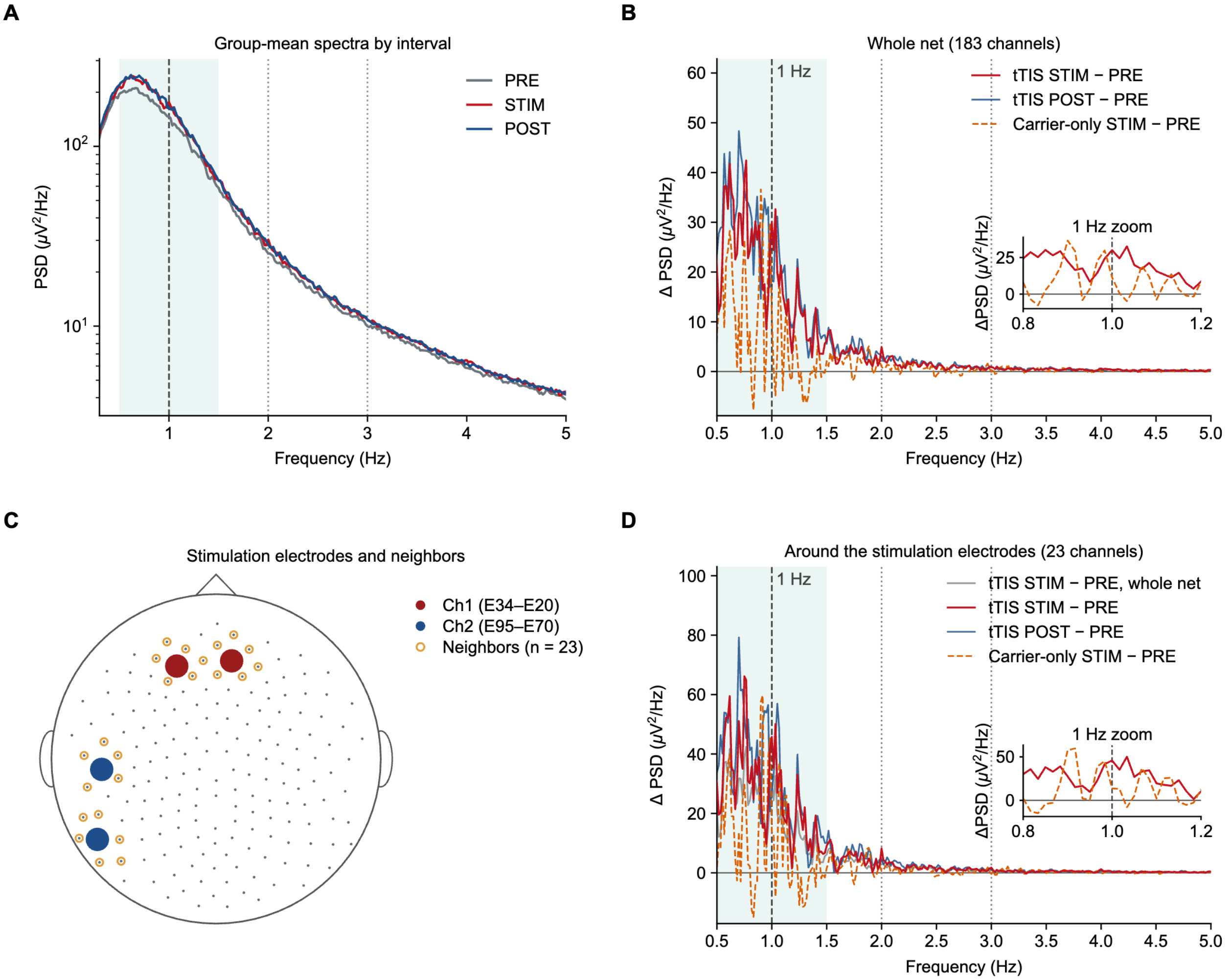


**Fig. S2.** *Spectral analysis. (A) Group-mean PRE, STIM, and POST power spectra for tTIS (n = 32; 60 s Welch windows, 0.0167 Hz resolution). Shading marks 0.5–1.5 Hz; markers indicate the 1 Hz envelope and 2/3 Hz harmonics. (B) Whole-net (183-channel) tTIS STIM-PRE and POST-PRE difference spectra, overlaid with the carrier-only STIM-PRE no-beat reference (n = 11; matched 15,000 Hz carriers). The inset enlarges 0.8–1.2 Hz. (C) The 183-channel net with the four stimulation electrodes (Ch1 E34+/E20−, Ch2 E70+/E95−) and their 23 first-ring neighbors in the sensor adjacency used for cluster tests. (D) As B, for the mean of the 23 neighbor channels only; the whole-net tTIS STIM-PRE difference is shown in gray for reference. Neither arm shows a narrow 1 Hz line in either channel set. All spectra are shown at full frequency resolution without smoothing.*

### Participant-scaled slow-wave detection

Each participant's YASA amplitude bounds were scaled by $k_i$, set from PRE negative half-waves alone (Methods). Fig. S3 shows the steps for one channel and the resulting bounds and detections in participants at the 10th percentile, median and 90th percentile of the cohort's amplitude scale.

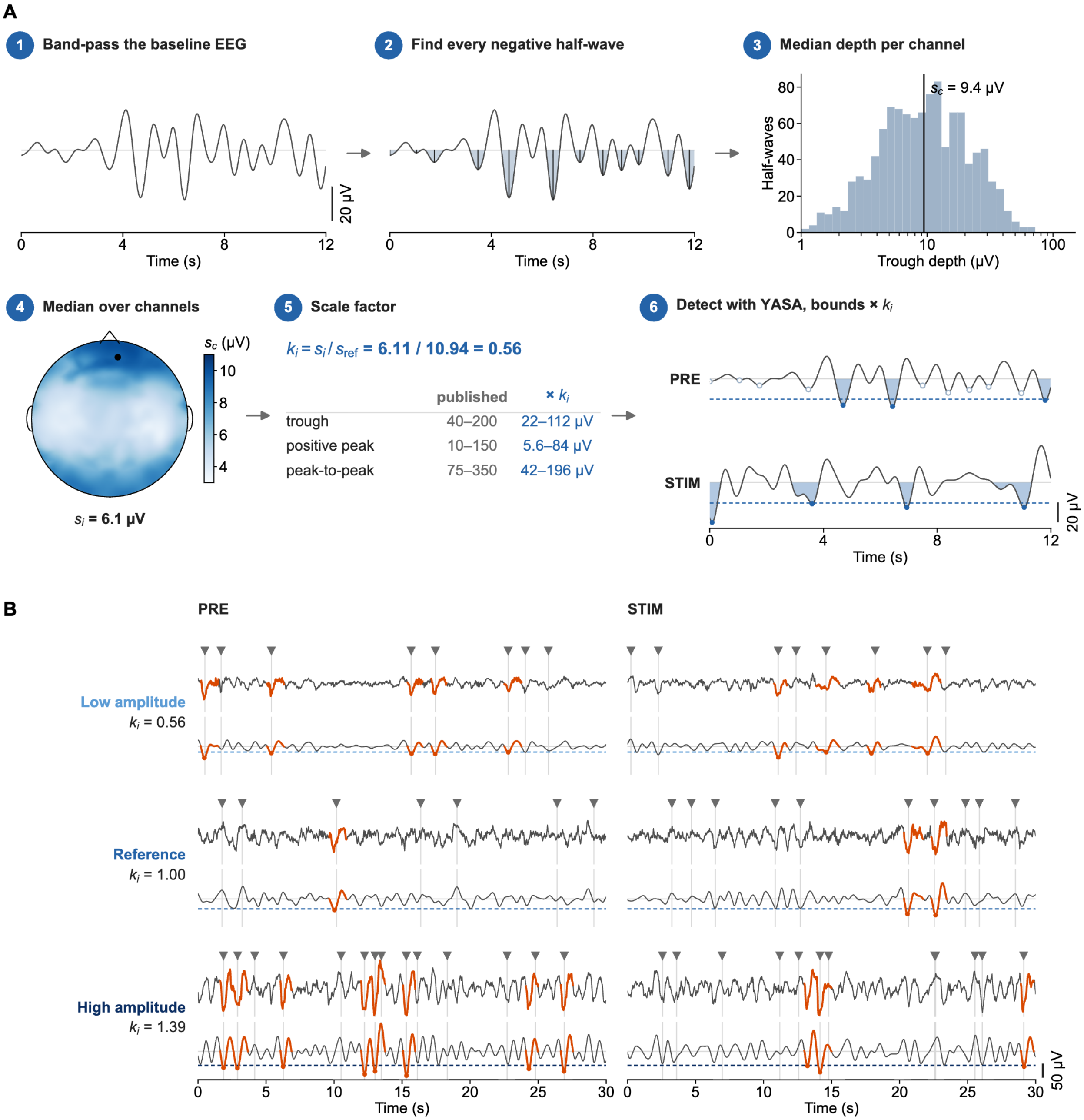


**Fig. S3.** *Participant-scaled slow-wave detection. (A) The steps for one tTIS participant and one frontal channel (E20). Steps 1–5 use PRE only, so the scale is blind to stimulation. (1) PRE EEG (average reference) band-passed at 0.3–1.5 Hz, as in YASA. (2) Every negative half-wave of 0.3–1.5 s is a candidate (shaded), with no amplitude rule, and its depth is how far its trough lies below zero. (3) The channel scale $s_c$ is the median depth of the channel's PRE half-waves. (4) The participant scale $s_i$ is the median of $s_c$ over the 183 channels (dot, E20). (5) The scale factor $k_i = s_i/s_{ref}$ multiplies every YASA amplitude bound. $s_{ref}$ = 10.94 µV is the cohort median, so a typical participant keeps the published bounds, and the duration criteria are unchanged. (6) YASA detection with the scaled bounds in PRE and STIM (N3). Dashed line, trough bound. Filled dots, detected troughs. Open dots, candidates that fail the scaled bounds. (B) Channel E20 in three tTIS participants at the 10th percentile, median and 90th percentile of $s_i$, over 30 s of PRE and STIM in N3. For each participant, the upper trace is the cleaned EEG and the lower trace the 0.3–1.5 Hz signal on which the bounds act (dashed line, trough bound). Waves detected on this channel are in orange, and triangles mark the participant's waves after merging across channels (Methods).*

### Cluster-based permutation statistics

The cluster-based permutation procedure common to the sensor- and source-space analyses is illustrated in Fig. S4.

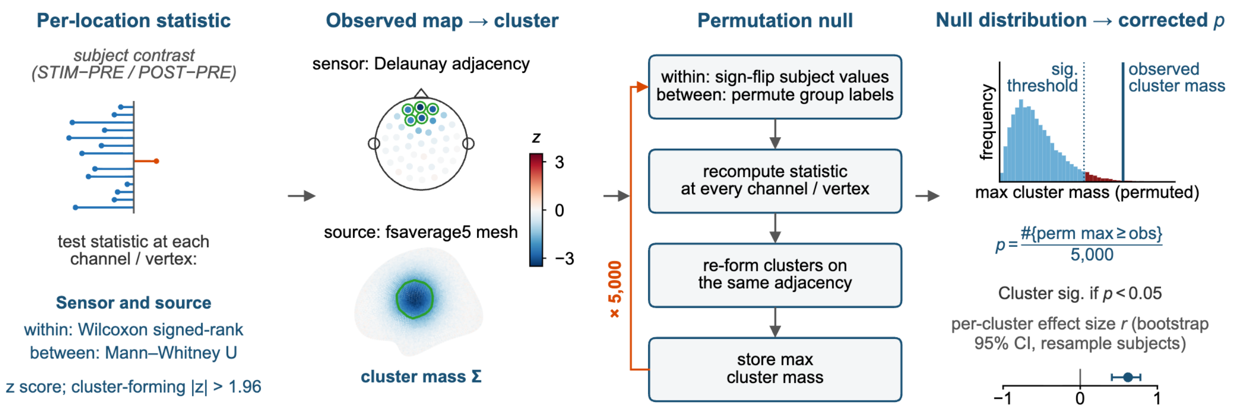


**Fig. S4.** *Cluster-based permutation statistics. Cluster-based permutation statistics, common to the sensor- and source-space analyses: a per-location rank statistic (within-group Wilcoxon signed-rank on paired changes, between-group Mann–Whitney U on change scores, at sensor channels and source vertices alike) is thresholded into candidate clusters on the sensor or source adjacency, and cluster mass is compared against a permutation null to yield cluster p-values. Values drawn are illustrative.*

### Sensor-space findings and within-night changes

**Table S3.** *Sensor-space cluster tests of slow-wave density and morphology.*

| Metric | Comparison | STIM/PRE: cluster p (q); channels | STIM/PRE: within-cluster r [95% CI]; p (q) | STIM/PRE: trend r; p | POST/PRE: cluster p (q); channels | POST/PRE: within-cluster r [95% CI]; p (q) | POST/PRE: trend r; p |
|---|---|---|---|---|---|---|---|
| Slow-wave density | tTIS | 0.0002 (0.0036); 174 | 0.85 [0.64, 0.98]; 0.000004 (0.00008) | −0.46; 0.023 | 0.0002 (0.0036); 176 | 0.83 [0.59, 0.98]; 0.000008 (0.00009) | −0.57; 0.0041 |
| Slow-wave density | Carrier-only | 0.95 (0.95) | 0.12 [−0.58, 0.76]; 0.76 (0.83) | 0.02; 1.0 | 0.21 (0.53) | 0.52 [−0.15, 0.91]; 0.15 (0.30) | −0.53; 0.16 |
| Slow-wave density | tTIS vs carrier-only | 0.030 (0.18); 30 | — | — | 0.23 (0.53) | — | — |
| Negative-peak amplitude | tTIS | 0.098 (0.35) | −0.53 [−0.83, −0.17]; 0.0083 (0.040) | 0.05; 0.80 | 0.16 (0.49) | −0.35 [−0.70, 0.02]; 0.084 (0.22) | 0.12; 0.56 |
| Negative-peak amplitude | Carrier-only | 0.61 (0.76) | 0.09 [−0.64, 0.70]; 0.83 (0.87) | −0.85; 0.014 | 0.80 (0.90) | −0.12 [−0.70, 0.61]; 0.76 (0.83) | −0.09; 0.85 |
| Negative-peak amplitude | tTIS vs carrier-only | 0.18 (0.49) | — | — | 0.88 (0.91) | — | — |
| Positive-peak amplitude | tTIS | 0.24 (0.53) | 0.28 [−0.11, 0.67]; 0.18 (0.33) | −0.10; 0.62 | 0.33 (0.59) | 0.30 [−0.10, 0.65]; 0.15 (0.30) | −0.00; 0.99 |
| Positive-peak amplitude | Carrier-only | 0.84 (0.91) | −0.36 [−1.00, 0.36]; 0.32 (0.51) | −0.13; 0.77 | 0.57 (0.76) | 0.15 [−0.58, 0.76]; 0.70 (0.83) | −0.45; 0.23 |
| Positive-peak amplitude | tTIS vs carrier-only | 0.39 (0.64) | — | — | 0.60 (0.76) | — | — |
| Peak-to-peak amplitude | tTIS | 0.17 (0.49) | 0.41 [0.02, 0.77]; 0.041 (0.15) | 0.08; 0.72 | 0.080 (0.32) | 0.41 [0.02, 0.75]; 0.043 (0.15) | −0.13; 0.54 |
| Peak-to-peak amplitude | Carrier-only | 0.86 (0.91) | −0.21 [−0.85, 0.45]; 0.58 (0.73) | 0.13; 0.77 | 0.71 (0.82) | 0.42 [−0.24, 0.91]; 0.24 (0.41) | −0.24; 0.56 |
| Peak-to-peak amplitude | tTIS vs carrier-only | 0.52 (0.72) | — | — | 0.69 (0.82) | — | — |
| Slope | tTIS | 0.51 (0.72) | 0.17 [−0.22, 0.58]; 0.41 (0.58) | 0.20; 0.32 | 0.27 (0.57) | 0.19 [−0.20, 0.58]; 0.35 (0.52) | 0.11; 0.60 |
| Slope | Carrier-only | 0.44 (0.69) | −0.61 [−1.00, 0.00]; 0.083 (0.22) | 0.35; 0.38 | 0.50 (0.72) | −0.97 [−1.00, −0.82]; 0.0020 (0.012) | 0.09; 0.85 |
| Slope | tTIS vs carrier-only | 0.30 (0.58) | — | — | 0.30 (0.58) | — | — |
| Frequency | tTIS | 0.080 (0.32) | −0.03 [−0.43, 0.36]; 0.88 (0.88) | 0.01; 0.98 | 0.022 (0.16); 13 | 0.13 [−0.26, 0.52]; 0.52 (0.70) | 0.10; 0.62 |
| Frequency | Carrier-only | 0.38 (0.64) | −0.52 [−0.94, 0.12]; 0.15 (0.30) | 0.09; 0.85 | 0.016 (0.15); 11 | −0.97 [−1.00, −0.82]; 0.0020 (0.012) | −0.09; 0.85 |
| Frequency | tTIS vs carrier-only | 0.051 (0.26) | — | — | 0.0008 (0.0096); 25 | — | — |

*Note. Each cell gives p with its Benjamini–Hochberg q in parentheses. Cluster p is each map's smallest cluster p (5,000 permutations; two-tailed family-wise error control within the map), q is across the 36 sensor maps, and the number after it counts the channels in clusters with cluster p < 0.05. Within-cluster r is the effect size of the paired change over the tTIS STIM/PRE density channels (subject-level bootstrap 95% CI), with its Wilcoxon signed-rank p (q across the 24 within-cluster tests; Fig. 3D). Trend gives the rank-biserial r of participants' within-night slopes across episodes and its Wilcoxon signed-rank p, which is uncorrected (Fig. 3A,B; Fig. S6).*

### Detectable effect sizes

With α = 0.05 (two-sided), the tTIS arm (n = 32) had 80% power to detect within-group paired effects of $d_z \geq 0.51$, the carrier-only arm (n = 11) only effects of $d_z \geq 0.94$, and the between-arm comparison only effects of $d \geq 1.00$; the carrier-only and between-arm null results therefore cannot exclude moderate effects. These figures assume a t test on one summary value per participant, not the permutation tests used for inference.

To estimate power with the permutation test itself, we drew 11 of the 32 tTIS participants at random 1,000 times and reran the within-group sensor density cluster test on each subset with the same adjacency, cluster-forming threshold and cluster mass, enumerating the sign-flip null exactly ($2^{11}$ = 2,048 permutations; Fig. S5). At the carrier-only sample size, a significant cluster ($p < 0.05$) was found in 82.4% of draws for STIM/PRE and 84.7% for POST/PRE, whereas the carrier-only arm's own test with the same exact null gave p = 0.95 and 0.22 (0.95 and 0.21 with the 5,000 random sign flips of the main analysis). Within the full-sample tTIS cluster, the mean density change of the drawn subsets (median 0.29 and 0.42 waves/min) exceeded the carrier-only change (0.04 and 0.22 waves/min) in all 1,000 draws for STIM/PRE and in 94.1% for POST/PRE. The detected tTIS increase alongside a null carrier-only result is therefore not explained by sample size alone, during or after stimulation.

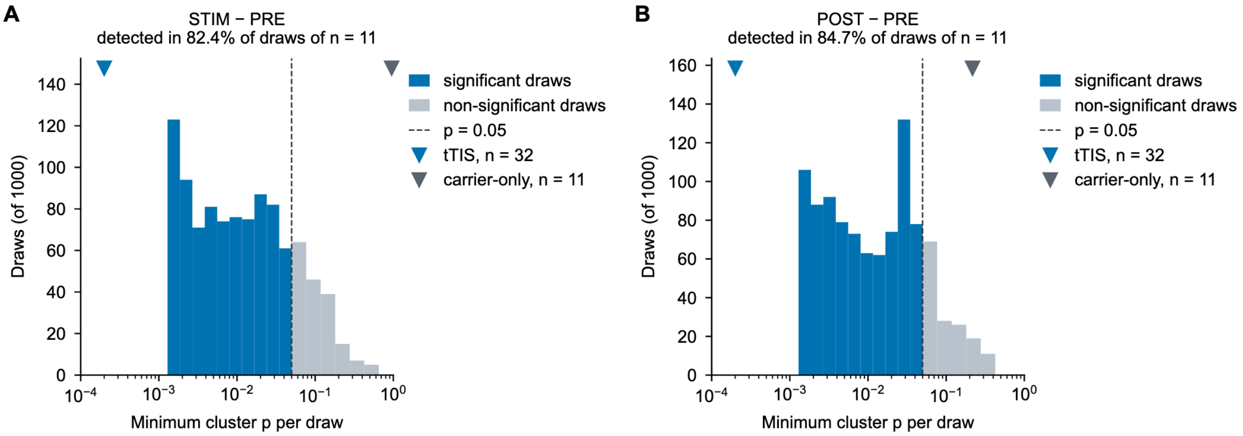


**Fig. S5.** *Sample-size-matched subsampling power of the within-tTIS density cluster test. Distribution of the minimum cluster p across 1,000 random draws of 11 of the 32 tTIS participants, each tested with the within-group sensor cluster permutation test used for inference (exact null, 2,048 sign flips). (A) STIM − PRE and (B) POST − PRE slow-wave density. Blue bars, draws with a significant cluster (p < 0.05, dashed line); gray bars, draws without one (every draw formed at least one cluster). Panel titles give the detection rate, the empirical power at the carrier-only sample size (82.4% for STIM/PRE and 84.7% for POST/PRE). Triangles mark the full tTIS test (n = 32, blue) and the carrier-only test (n = 11, gray, exact null; p = 0.95 and 0.22).*

### Time-of-night dependence of wave morphology

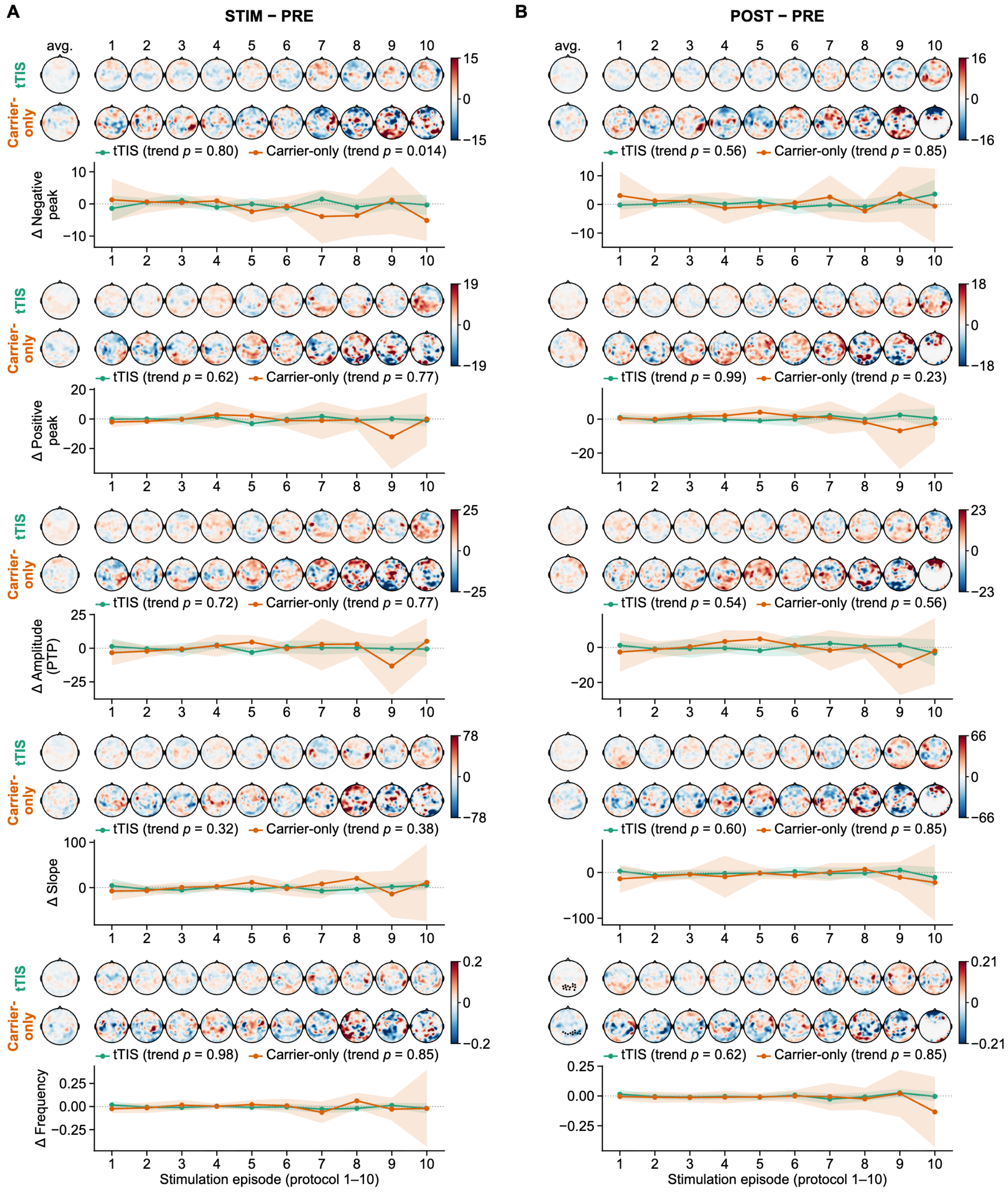


**Fig. S6.** *Time-of-night morphology (tTIS $n = 32$, carrier-only $n = 11$). Left, pooled channel-wise changes across episodes in negative-peak, positive-peak and peak-to-peak amplitude, slope and frequency, with tTIS above carrier-only and channels in significant clusters dotted (corrected $p < 0.05$ within the map; Table S3). Only frequency after stimulation formed clusters, in tTIS (13 channels) and carrier-only (11 channels). Right, per-episode topographies above the within-night trend across the first ten stimulation episodes. Points show the group-mean change at each episode (± subject-level 95% CI); $p$ is the within-night trend test (Wilcoxon signed-rank test of participants' slopes across episodes), given for each group in the legend (carrier-only $n = 10$ with ≥ 4 episodes). Pooled and per-episode maps share a symmetric color scale within each metric and contrast. Blank areas in late carrier-only episode maps are channels where no participant had waves in both segments. Amplitudes are in μV on each participant's own recording and are not comparable between participants, because detection bounds scale with $k_i$.*

### Within-night decline: protocol order versus cleaned-recording time

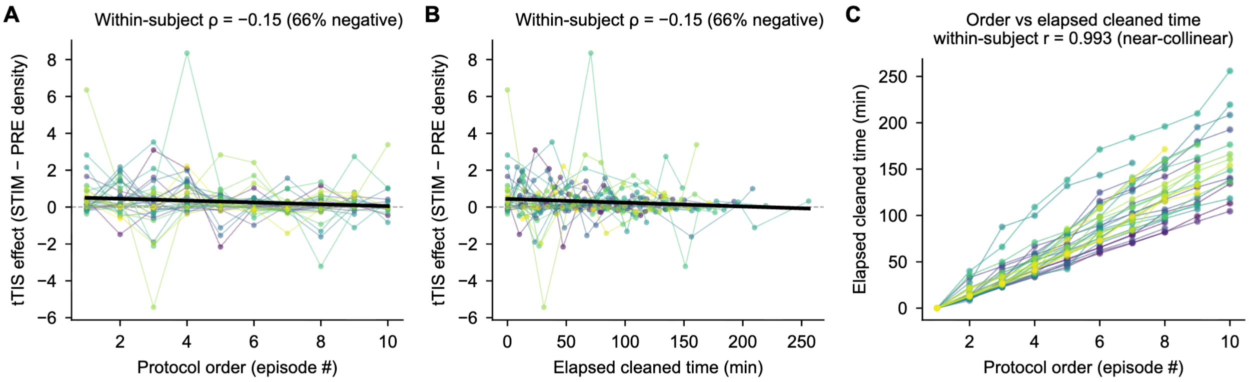


**Fig. S7.** *Within-night tTIS effect by protocol order and elapsed cleaned-recording time (32 participants; first ten protocols, 292 episodes). Channel-mean STIM-minus-PRE density versus (A) episode number and (B) elapsed cleaned-recording time in minutes, derived from each protocol's earliest detected-wave start relative to the earliest across protocols (thin lines, participants; bold line, group linear fit, −0.050 waves/min per episode). Because elapsed time rises monotonically with episode number within each participant, both axes give the same mean within-subject Spearman ρ (−0.15, negative in 66% of participants). (C) The axes are near-collinear within participants (mean Pearson r = 0.993).*

### Source-space findings

**Table S4.** *Source-space cluster tests of slow-wave origin and involvement.*

| Map | Comparison | STIM/PRE: cluster p (q); significant clusters (vertices) | POST/PRE: cluster p (q); significant clusters (vertices) |
|---|---|---|---|
| Origin, probability (P) | tTIS | 0.83 (0.83) | 0.32 (0.42) |
| Origin, probability (P) | Carrier-only | 0.065 (0.11) | 0.83 (0.83) |
| Origin, probability (P) | tTIS vs carrier-only | 0.59 (0.70) | 0.027 (0.065); 1 (166) |
| Involvement, probability (P) | tTIS | 0.0036 (0.014); 2 (7,487) | 0.0028 (0.014); 2 (12,405) |
| Involvement, probability (P) | Carrier-only | 0.011 (0.032); 1 (2,616) | 0.0024 (0.014); 2 (7,973) |
| Involvement, probability (P) | tTIS vs carrier-only | 0.067 (0.11) | 0.10 (0.15) |

*Note. Each cell gives the map's smallest cluster p (family-wise error controlled within the map; 5,000 permutations) with its Benjamini–Hochberg q across the 12 source maps in parentheses, followed by the significant clusters. P, per-wave probability.*

### Source origin and involvement probability maps

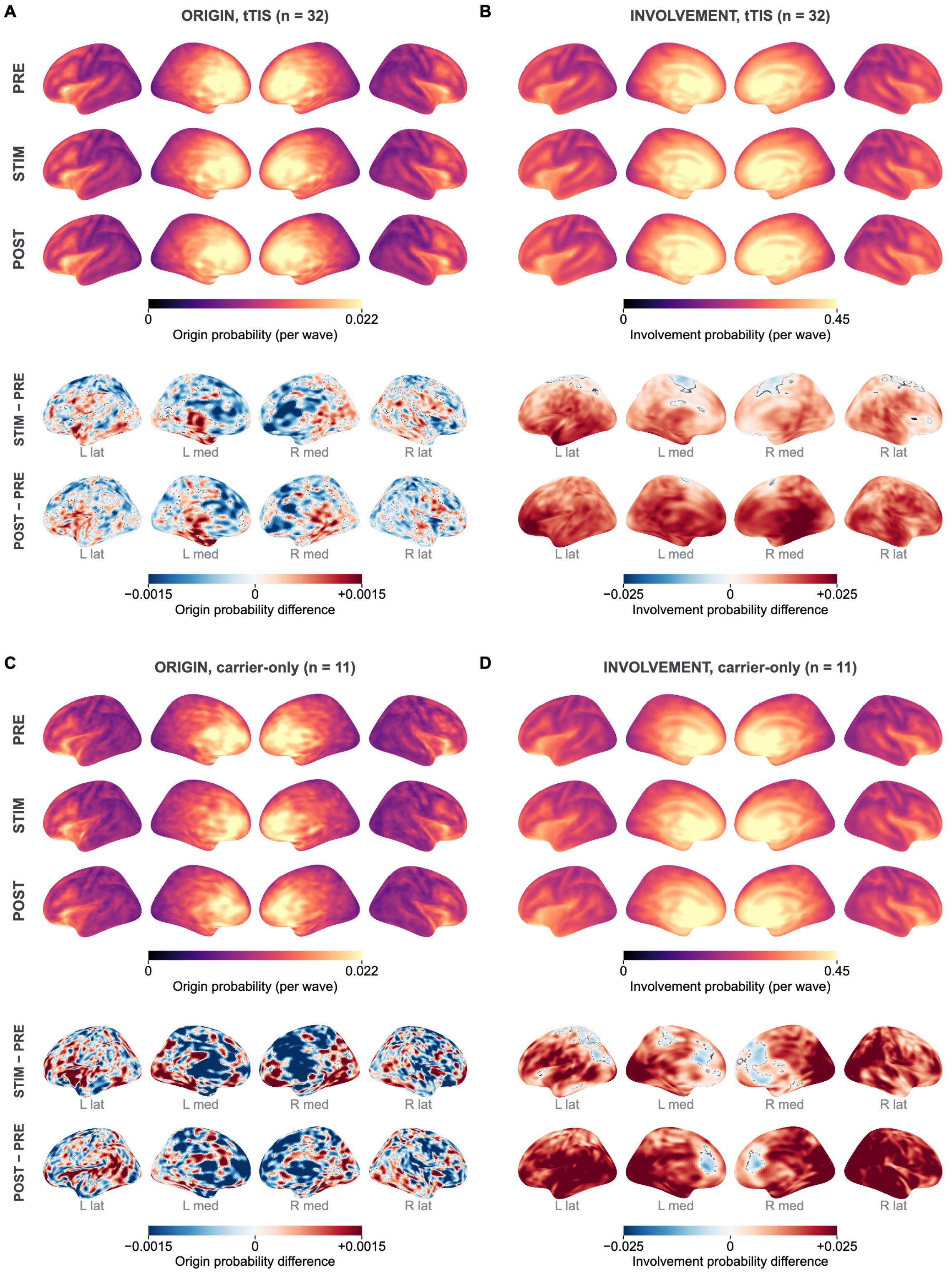


**Fig. S8.** *Slow-wave source probabilities (P) for origin and involvement in tTIS (A, B; n = 32) and carrier-only (C, D; n = 11) in PRE, STIM and POST, shown on inflated fsaverage5 (LH lateral, LH medial, RH medial, RH lateral). Origin, the earliest 10% of vertices to reach their local maximum of relative current ; involvement, current amplitude at the negative peak above the vertex's own 95th percentile at random PRE times. Vertex values are the share of a participant's waves that originate at or involve the vertex, averaged across each arm's participants. Color limits are the 98th percentile of the pooled tTIS values (origin 0.022, involvement 0.45) and of the pooled absolute tTIS differences (origin 0.0015, involvement 0.025), and both arms share them, so carrier-only differences beyond them saturate. Lower rows show descriptive differences. Involvement probability rose in both arms during and after stimulation (Fig. 4), whereas origin probability showed no significant change (Table S4).*

### Participant time courses of the shift toward N3 sleep

The time course of Fig. 6A splits the tTIS group at the median whole-cortex $TI_{max}$, while the test uses the continuous field. Fig. S9 shows every participant's change in N3 share at every step, ordered by the whole-cortex field.

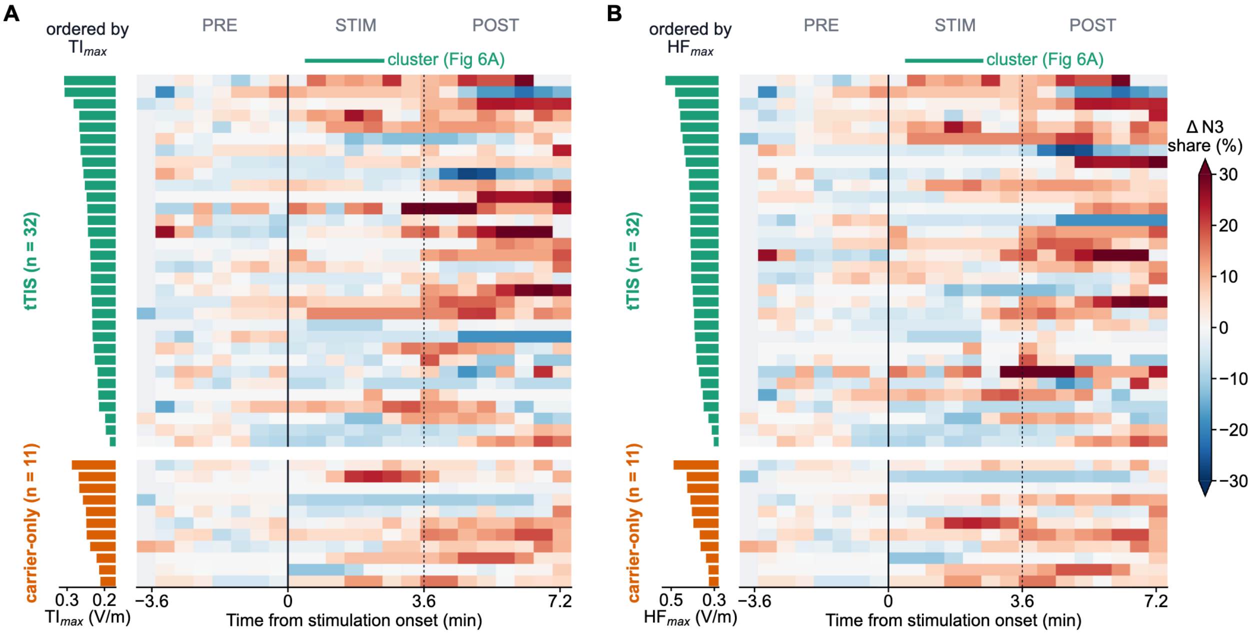


**Fig. S9.** *Change in N3 share for every participant, ordered by modeled field. Each row is one participant and each column one 30-s step aligned to stimulation onset; color is the participant's change in N3 share (% of N2 + N3) from the participant's PRE mean, pooled over all of the participant's stimulations. Rows are ordered by whole-cortex $TI_{max}$ (A) or $HF_{max}$ (B), shown as bars at left, tTIS above and carrier-only below (carrier-only $TI_{max}$ is the placebo modulation depth). The bar above each map is that field's cluster of steps in which the change followed the field across the tTIS participants (30 to 150 s; Fig. 6A); the dotted line marks the median end of STIM, and light gray cells are steps a participant does not reach. Step position is the order of the usable 10-s bins in their window × 10 s, so it follows clock time to within about 20 s.*

### Leave-one-subject-out sensitivity

**Table S5.** *Leave-one-subject-out sensitivity across reported analyses.*

| Analysis | Contrast | Omitted | LOSO result |
|---|---|---|---|
| Sensor density | STIM/PRE | 32 tTIS | Cluster $p < 0.05$ in 32/32; worst $p = 0.0002$; max $\|\Delta r\| = 0.07$ |
| Sensor density | POST/PRE | 32 tTIS | Cluster $p < 0.05$ in 32/32; worst $p = 0.0002$; max $\|\Delta r\| = 0.08$ |
| Sensor density, tTIS − carrier-only | STIM/PRE | 43 (32 tTIS, 11 carrier-only) | Cluster $p < 0.05$ in 24/43; worst $p = 0.19$; max $\|\Delta r\| = 0.11$ |
| Sensor negative-peak amplitude | STIM/PRE | 32 tTIS | $r = -0.62$ to $-0.50$; max $\|\Delta r\| = 0.09$; Wilcoxon $p < 0.05$ in 32/32 |
| Sensor peak-to-peak amplitude | STIM/PRE | 32 tTIS | $r = 0.38$–$0.50$; max $\|\Delta r\| = 0.09$; Wilcoxon $p < 0.05$ in 11/32 |
| Sensor peak-to-peak amplitude | POST/PRE | 32 tTIS | $r = 0.37$–$0.49$; max $\|\Delta r\| = 0.08$; Wilcoxon $p < 0.05$ in 12/32 |
| Sensor slope, carrier-only | POST/PRE | 11 carrier-only | $r = -1.00$ to $-0.96$; max $\|\Delta r\| = 0.03$; Wilcoxon $p < 0.05$ in 11/11 |
| Sensor frequency, carrier-only | POST/PRE | 11 carrier-only | $r = -1.00$ to $-0.96$; max $\|\Delta r\| = 0.03$; Wilcoxon $p < 0.05$ in 11/11 |
| Source involvement P | STIM/PRE | 32 tTIS | $r = 0.70$–$0.82$; max $\|\Delta r\| = 0.10$ |
| Source involvement P | POST/PRE | 32 tTIS | $r = 0.76$–$0.83$; max $\|\Delta r\| = 0.05$ |
| Source involvement P, carrier-only | STIM/PRE | 11 carrier-only | $r = 1.00$–$1.00$; max $\|\Delta r\| = 0.00$ |
| Source involvement P, carrier-only | POST/PRE | 11 carrier-only | $r = 1.00$–$1.00$; max $\|\Delta r\| = 0.00$ |

*Note. Sensor clusters were retested with 5,000 permutations per omission. Morphology rows use the tTIS STIM/PRE density-significant channels and the nominal Wilcoxon p. Source effects use the smallest-p full-sample cluster. Source spatial significance was not retested, and origin probability, with no significant full-sample cluster, was not retested. Carrier-only involvement rows are conditional on clusters found in the same 11 participants. Fixed-region analyses assess participant sensitivity, not independent spatial validation.*

### Supplementary References

**1.** Franke, M., Bhadra, N., Bhadra, N. & Kilgore, K. Direct current contamination of kilohertz frequency alternating current waveforms. *Journal of Neuroscience Methods* **232**, 74–83 (2014).

**2.** von Conta, J., Kasten, F. H., Ćurčić-Blake, B., Schellhorn, K. & Herrmann, C. S. Characterizing low-frequency artifacts during transcranial temporal interference stimulation (tTIS). *Neuroimage: Reports* **2**, 100113 (2022).
